# Evolutionary implications of phenotype switching for growth and survival of tumors in stochastic environments

**DOI:** 10.64898/2026.08.18.745434

**Authors:** Sanasar G. Babajanyan, Yuri I. Wolf, Rafael R. Canevarolo, Praneeth R. Sudalagunta, Ariosto S. Silva, Eugene V. Koonin, Erez Persi

## Abstract

Tumors evolve under constantly changing microenvironmental conditions, and a key major mechanism by which cancer cells adapt to fluctuating environments is stochastic phenotypic switch via epithelial–mesenchymal plasticity (EMP). Here we demonstrate the existence of EMP in human breast epithelial cell lines and assess the fitness of E and M cells in environments that fluctuate between favorable and harsh conditions. We then develop a theoretical framework for analysis of tumor evolution with E-M stochastic phenotypic switch that integrates the behaviors of E and M cells. We characterize the outcomes of tumor evolution via three characteristic times describing i) tumor growth, ii) phenotypic adaptation and iii) fluctuations of the environment. We identify a tradeoff between tumor growth and survival, with different phenotype switching rates maximizing each of these objectives. An anticancer therapeutic strategy is proposed based on tumor survival analysis, revealing that blocking mesenchymal to epithelial transition, rather than epithelial to mesenchymal transition, is critical under fluctuating conditions. Thus, this study elucidates fundamental evolutionary mechanisms of tumor phenotypic adaptation in fluctuating environments with potentially important clinical implications.

**SIGNIFICANCE:** Tumors evolve under constantly changing environmental conditions, to which they adapt primarily via epithelial-mesenchymal plasticity. However, the impact of fluctuating environment on tumor growth and phenotypic composition remains elusive. We developed a mathematical model of tumor growth under fluctuating conditions with stochastic phenotypic epithelial-mesenchymal switch as the main mechanism of adaptation. Motivated by experimentally observed differences in the growth of epithelial and mesenchymal cells under variable conditions, the model reveals non-trivial evolutionary outcomes, depending on the relative time scales of the underlying processes, reminiscent of the Parrondo’s Paradox in game theory. Based on the model analysis, an optimal therapeutic strategy is proposed, with the mesenchymal-epithelial transition identified as the critical target.

## INTRODUCTION

Both organisms and tumors typically evolve in environments with fluctuating parameters such as nutrient availability, temperature, and pH. Fluctuating environments impose variable selective pressures to which evolving populations respond and adapt using diverse genetic and non-genetic mechanisms, including population-level diversity as well as cellular phenotypic plasticity, depending on the time scale and nature of environmental fluctuations [1–3]. Environmental fluctuations, both deterministic and stochastic, can lead to evolutionary outcomes that differ drastically from those in the corresponding fixed environments [4–9].

In unpredictable environmental conditions, genetic variation can reduce the overall fitness of a population, lead to accumulation of deleterious mutations and increase the probability of extinction [10–14]. In this context, the optimal adaptive strategy is not a cue-based response but rather stochastic phenotypic switching, whereby populations exhibit alternative phenotypic states, maximizing the chance that some individuals will cope better with future unknown conditions [15–19].

Both theory and experimental evidence suggest that the rate of such stochastic switching is tuned to the time scale of environmental fluctuations, maximizing the long-term fitness of the population in temporally uncorrelated and exponential growth phases [4, 20–22]. However, maximizing the long-term growth is not the safest path for a population to evolve because fluctuations can cause population extinction [23, 24]. Therefore, optimizing long-term growth and maximizing survival time might select distinct biological traits.

Similarly to species evolution, cancers also evolve and adapt to dynamic and heterogeneous microenvironmental selection pressures, imposed by nutrient and oxygen availability, immune surveillance and therapy, which leads to the emergence of highly plastic, generalist and invasive phenotypes [25–27]. A central mechanism of such adaptation is the epithelial-to-mesenchymal transition (EMT), a reversible program conferring invasiveness, resistance to apoptosis, and stem-like properties, the reversal of which, the mesenchymal-to-epithelial transition (MET), facilitates metastatic colonization at distant sites [28–33].

However, in contrast to a fixed environment, the population-level consequences of EMT/MET switching under temporally correlated fluctuations of microenvironmental conditions remain largely unexplored, [34, 35].

Another complication in tackling cancer growth is the limited availability of resources in the microenvironment, because of which the long-term fitness of cancer, that is, the logarithmic population size, remains coupled to the current population size and composition of the cancer due to phenotypic heterogeneity [4, 22]. Thus, it remains unknown how the characteristics of environmental fluctuations determine the optimal EMT/MET switching rate and impact the growth of subpopulations, and what is the optimal therapeutic strategy to compact tumor growth under such conditions.

Here, we first explore experimentally how epithelial, *E*, and mesenchymal, *M*, cells respond to environmental fluctuations and then develop a population genetics theory incorporating the dynamics of EMT/MET. We consider the evolutionary outcome of cancer growth in a stochastic environment, with finite correlation times between switches, that is, under colored environmental noise [36, 37]. We derive the evolutionary outcomes for tumor size and compositional variability as functions of phenotypic switching rates and statistics of environmental fluctuations. The model allows us to define the optimal EMT/MET switching rates that provide the maximal long-term growth rate or minimal extinction risk for the cancer. By considering the extinction risk, we identify the functional relationship between the switching rates, environmental bias and selection strength that results in the extinction of tumor. Our results suggest that inhibiting MET, rather than EMT, would result in more effective elimination of tumors, providing actionable insights for evolutionary-informed therapeutic intervention aimed at suppressing cancer progression.

## RESULTS

### Experimental motivation: MCF10A-derived tumorigenic clones exhibit a continuum of epithelial–mesenchymal states

Epithelial–mesenchymal plasticity (EMP), encompassing interconversion of the *E* and *M* cellular states through epithelial–mesenchymal transition (EMT) and mesenchymal–epithelial transition (MET), plays a central role in cancer progression, metastasis, and therapeutic resistance. Despite its biological importance, relatively few in vitro models spontaneously recapitulate the phenotypic diversity and dynamic plasticity associated with EMP. Here, we describe an EMP model that evolved from the non-tumorigenic breast epithelial cell line MCF10A following approximately two years of selection under cyclical nutrient-depletion stress. The resulting cell populations recapitulate several features associated with EMP in breast cancer, including the emergence of distinct *E* and *M* phenotypes with dynamic transitions between these states, and the spontaneous formation of multicellular spheroids containing both *E* and *M* cells. We previously reported that long-term selection of MCF10A cells under cyclical nutrient depletion generated tumorigenic clones, with acquired driver mutations and increased ROS and lactate production relative to parental cells, indicating mutagenesis and acquisition of the Warburg phenotype [38]. Comprehensive molecular characterization of representative clones using whole-exome sequencing (WES), single-cell multiome profiling (sc-Multiome; paired single-cell RNA/ATAC-seq), and copy-number variation analysis revealed substantial genotypic diversity among these clones despite their common ancestral origin. Despite this genetic divergence, we observed phenotypic convergence to EMP, whereby clones differed by the ratio of cells in each of these two states. All clones dynamically switch between *E* and *M* states, mediated by epigenetic activation of Grainyhead-like 2 transcription factor (GRHL2), a master regulator of epithelial identity and the associated transcriptional and epigenetic programs [38] consistent with previous studies ([39, 40]). Under standard culture conditions, the *E*-dominant clones PE14 and PE16 retained strong epithelial morphology and transcriptional state, whereas PE26 adopted a highly mesenchymal phenotype, and PE21 exhibited intermediate characteristics. Together, these evolutionarily related but phenotypically divergent populations provide an experimental framework for investigating the mechanisms governing EMP and the influence of environmental conditions on transitions between cellular states. Here, we use these clones to further investigate how environmental stress affects the balance between *E* and *M* states. Under conditions of continuous media replacement, all clones rapidly expanded and reached full confluency (Fig.1A-B, and SI section I Supplementary Figure 2.A-C). However, important differences emerged after confluency: the *E*-dominant clones consisted of a largely epithelial monolayer with localized populations of *M*-like cells growing above the epithelial layer. In contrast, the *M*-dominant PE26 clone continued to proliferate beyond confluency and progressively formed large three-dimensional spheroid-like structures that grew on top of the mesenchymal monolayer (SI section I Supplementary Figure 2C,I). Previous immunofluorescence analyses demonstrated that these spheroids exhibit distinct spatial organization, with *E*-like cells enriched in the core and *M*-like cells preferentially localized at the periphery of the spheroids [38]. These observations suggest that maintenance of optimal environmental conditions facilitate the expansion and spatial organization of *M* states, particularly in clones whose EMP balance favors *M* states. Next, clones were cultured without media replacement and monitored until loss of confluency occurred (Fig.1C-D, and SI section I Supplementary Figure 2D-F). All clones exhibited similar early growth dynamics: starting from approximately 25% confluency, cultures rapidly expanded and reached full confluency within three to six days. Representative brightfield images are shown in Fig.1E-F, and SI section I Supplementary Figure 2G-I: each clone reached initial confluency followed by cell death due to nutrient starvation, and a subsequent re-growth after media replenishment. Following initial confluency, we noticed a characteristic biphasic response in all clones during the starvation phase, marked by a rapid decline followed by an extended plateau. Visual inspection of sequential images confirmed that the first phase is characterized by the death of the *M* cells, whereas *E* cells survived through the second phase, suggesting that *M* cells are more sensitive to nutrient depletion than *E* cells. Although the *E*-dominant clones (PE14 and PE16), as well as parental cells, underwent a rapid population decline resulting in loss of nearly 40-50% cells from maximum confluency, the *M*-dominant clones (PE21 and PE26) show a more precipitous drop, with 75-100% of the cells dying. These observations agree with the proportion of cells in *M* and *E* states, as assessed by scMultiome, in these clones (only 36% and 20% cells in *E* state in PE21 and PE26, respectively, compared to 94% in the other clones [38]). These observations suggest that *M* populations benefit from optimal nutrient availability by growing beyond confluency, forming three dimensional structures, whereas *E* populations exhibited substantial resilience under prolonged nutrient depletion. This dichotomy is in part due to the *E*-state associated cell-cell adhesion signaling, which induces cell cycle arrest in high confluency conditions, and promotes survival under stress [29, 41–44]. These observations suggest the existence of fundamentally distinct ecological strategies among tumorigenic clones and provide the experimental foundation for the mathematical modeling framework developed in this study.

**FIG. 1:**
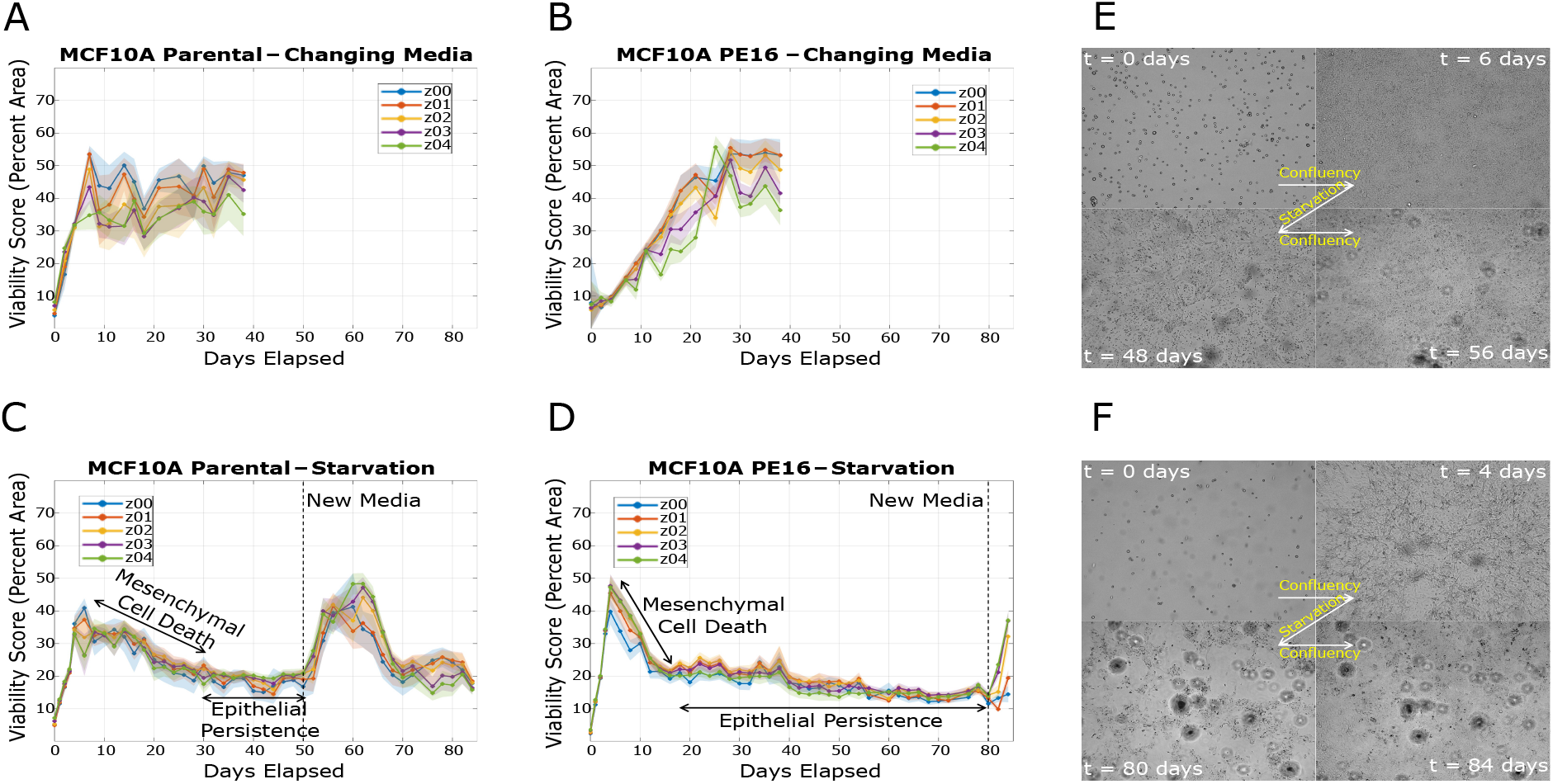
Nutrient deprivation induces a biphasic response in parental MCF10A and tumorigenic clones. **A)** and **B**) show cells maintained under media-changing conditions for MCF10A parental and *E*-dominant PE16 clone, respectively. Under these nutrient-replete conditions, areas occupied by live cells generally increased or remained stable, reflecting continued growth and maintenance of adherent cultures. **C)** and **D)** show matched starvation/re-feeding conditions for MCF10A parental and tumorigenic clones, respectively. In these cultures, cells initially reached high confluency, followed by a biphasic starvation response: an early decline in viability corresponding to the mesenchymal death phase, and a later plateau corresponding to persistence of an adherent *E*-like population. **E)** and **F)** show representative brightfield images illustrating the transition from initial seeding to confluency, starvation-associated loss of live cells, persistence of a subpopulation, and subsequent confluency upon media replenishment corresponding to the experimental conditions C) and D), respectively. Together, these data show that standard media replacement supports continued culture viability, whereas nutrient/media deprivation reveals clone-specific starvation sensitivity characterized by early cell death of M cells followed by long-term persistence of residual adherent cells.

### Mathematical model of tumor evolution in stochastic environment

Motivated by the above experimental observations, we explored a mathematical model of the growth of phenotypically heterogeneous solid tumors in temporally correlated stochastic environment. Under the model, the tumor consists of *E* and *M* cells that compete for shared resources limited by the carrying capacity of the environment, *K*. In addition to reproduction and competition, cells can reversibly switch between epithelial and mesenchymal phenotypic states.

Environmental fluctuations affect only the reproduction rate of mesenchymal cells, that is *r*_*M*_ = *r*_*E*_ + *ϵξ*(*t*) where *ξ*(*t*) = ± 1 is a colored dichotomous noise [36, 45]. The parameter *ϵ* quantifies the selective advantage or disadvantage of mesenchymal cells compared to epithelial ones. For simplicity, the growth rate of epithelial cells is fixed to *r*_*E*_ = 1. Environmental transitions occur with time-independent rate *λ*, while *δ* ∈ [− 1, 1] controls the bias toward favorable or unfavorable conditions for mesenchymal cells, so that the stationary probabilities of the good and bad environmental states are 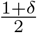 and 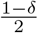, respectively. The mean reproduction rate of *M* cells, averaged over environmental fluctuations, is given by *E*_*ξ*_[*r*_*M*_] = 1 + *ϵδ*. For unbiased environment, that is *δ* = 0, *E*_*ξ*_[*r*_*M*_] = *r*_*E*_ = 1.

Let *x* ∈ [0, 1] denote the fraction of *E* cells in the tumor and *N* the total number of cells. The dynamics of these variables are governed by a piecewise-deterministic Markov process

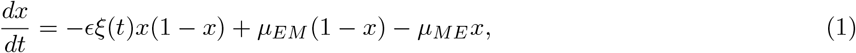

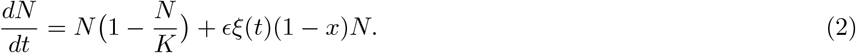

Here *µ*_*EM*_ and *µ*_*ME*_ are the phenotype-switching rates from mesenchymal to epithelial states and vice versa. Equations (1) and (2) are obtained from a two-species Lotka-Volterra competition model through change of variables (see SI section II). In the absence of environmental fluctuations, the tumor size in the stable equilibrium is equal to the carrying capacity of the environment, *K*.

The equilibrium fraction of epithelial cells is 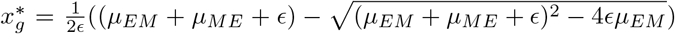 and 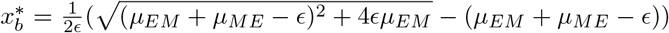 in good and bad environment, respectively. Because mesenchymal cells are selectively favored when *ξ* = 1 and disfavored when *ξ* = − 1, it follows that *x*.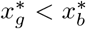 Thus, the tumor composition at equilibrium shifts toward mesenchymal dominance in good periods and toward epithelial dominance in bad periods of environmental fluctuations, respectively.

The size of the tumor population in these equilibrium states is given by 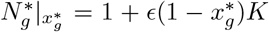 and 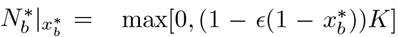 in good and in bad conditions, respectively. Due to the environmental switches, the long-term dynamics need not coincide with any equilibrium associated with a fixed environment, in general.

A key feature of the model is the asymmetric coupling between the tumor composition and population size. Equation (2) shows that tumor growth explicitly depends on the current fraction of epithelial cells in the tumor *x*, implying that phenotypic heterogeneity directly influences the tumor size dynamics. In contrast, under Eq. (1), *x* is independent of *N*, that is, the evolution of tumor composition is driven solely by environmental selection and phenotype switching.

### Time-scale analysis of tumor evolution

The evolutionary outcomes of tumor proliferation are governed by the interplay of three distinct processes: environmental switching, phenotypic adaptation, and tumor growth. Each process is characterized by its own intrinsic timescale, and the relative ordering of these timescales determines the long-term behavior of the system. Different timescale orderings give rise to qualitatively different stationary distributions of tumor composition and population size, which can be unimodal, bimodal or multimodal.

To characterize these evolutionary regimes, we first identify the characteristic timescales associated with each process. The tumor composition evolves on a time scale 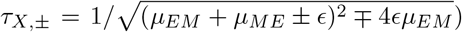 whereas the characteristic growth time of a tumor with a fixed composition *x* is *τ*_*N*,_^*±*^ = 1*/* |1 _*±*_ *ϵ*(1 − *x*) (see SI section II for details). Environmental fluctuations are characterized by the average persistence times of the good and bad states *τ*_*ξ*,_ ± = 1*/λ* ±= 1*/λ*(1 ∓*δ*).

The coupling between the population size of the tumor and the fraction of epithelial cells in the tumor does not allow us to find the exact joint probability distribution function of *N* and *x*. We approximate the dynamics of the tumor size, (4), by treating the fraction of epithelial cells as fixed. This approximation allows us to derive the stationary distributions for the tumor size and its composition.

Within this approximation the stationary probability density functions of tumor composition and population size for a fixed composition are as follows

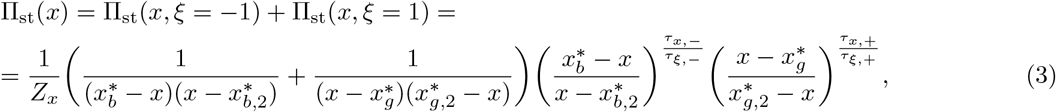

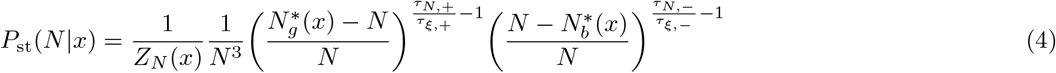

In (3), Π_st_(*x, ξ* = ± 1) denotes the joint stationary probability of observing tumor composition *x* while the environment is in state *ξ* = ± 1. *Z*_*x*_ is the normalization constant. The stationary probability density function involves non-physical equilibrium states of the tumor 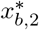 and 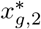 as well. These states lie outside of the physically admissible region 0 ≤*x* ≤1. Although these steady states are not realized dynamically, they define the shape of the distribution function.

In (4), the endpoints of the support correspond to equilibrium states of the tumor population size in a fixed environment for a given composition of the tumor, that is, 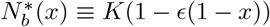 and 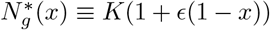 Note that the support of the tumor population size distribution for a given composition of the tumor, that is, 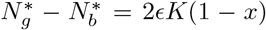, narrows with increasing fraction of *E* cells in the tumor. The normalization explicitly depends on the tumor composition, *Z*_*N*_ (*x*) in (4) (for further details on the derivation of the pdfs refer to SI section III).

The probability density functions (3) and (4) allow us to retrieve the information on the evolutionary outcomes of the tumor growth and phenotypic composition.

To illustrate the role of timescale ordering in evolutionary outcomes of the tumor proliferation, we focus on the unbiased environment, *δ* = 0, with symmetric phenotype switching rates, *µ*_*EM*_ = *µ*_*ME*_ = *µ*. In this case, the expressions for the characteristic timescales simplify to *τ*_*X,±*_ = *τ*_*X*_ = (4*µ*^2^ + *ϵ*^2^)^*−*1*/*2^ and *τ*_*ξ,±*_ = *λ*^*−*1^ for phenotypic adaptation and environmental fluctuation, respectively. Nevertheless, the analysis via characteristic timescales provided below holds for asymmetric phenotype switching rates and for biased environments as well (see SI section III for further details).

We first consider a fast-fluctuating regime, whereby environmental switching occurs faster than phenotypic adaptation, *τ*_*ξ*_ *< τ*_*X*_ . Here, the tumor composition does not relax toward the equilibrium states 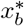 and 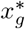 before the environment changes again, on average. Indeed, near the equilibrium state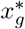 the stationary probability behaves as 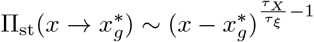 an analogous expression holds for 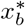 as well. Since *τ*_*ξ*_ *< τ*_*X*_ the stationary probability vanishes at both boundaries. Therefore, it is unlikely to observe the equilibrium states of tumor composition associate with fixed environments.

For sufficiently ffast environmental switching, such that 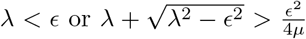 with *λ > ϵ*, the stationary distribution typically has a single peak around 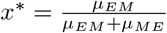 which reduces to *x*^*∗*^ = 1*/*2 for symmetric rates (see Fig.2A top left panel). *x*^*∗*^ corresponds to self-averaged noise limit whereby the noise can be substituted by its mean value. However, *x*^*∗*^ can become a local minimum of the probability function when 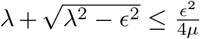 and *λ > ϵ*. In this case, the stationary probability functions become *M*-shaped, characterized by two symmetric maxima around *x*^*∗*^, (see Fig.2A middle left panel and SI section IV for further details) .

**FIG. 2:**
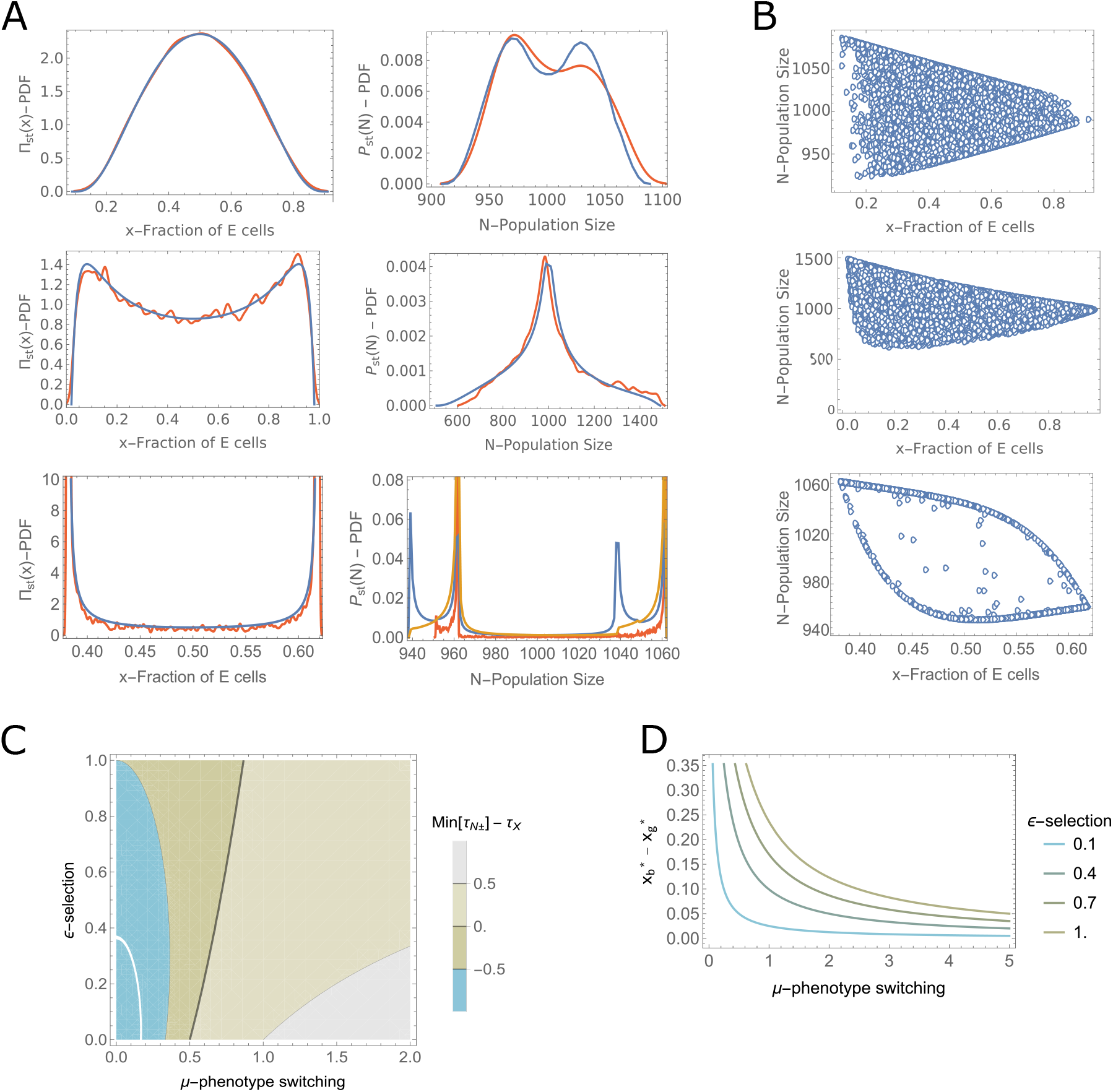
Evolutionary outcomes for different orderings of the characteristic timescales of tumor growth, phenotypic adaptation and environmental fluctuations. **A)** Probability density functions of tumor composition and population size (left and right in each row respectively). Blue and red lines in each panel show the pdf of the respective quantity obtained from the model, through (3), (4) and (5), and through simulations of stochastic equations (1) and (2), with 10^4^ independent runs with randomly chosen initial states. Fast environmental switching regime, whereby environment fluctuates faster compared to the phenotypic change of tumor but slower than tumor growth (*τ*_*X*_ *> τ*_*ξ*_ *> τ*_*N*_) with parameters *ϵ* = 0.1, *µ*_*EM*_ = *µ*_*ME*_ = 0.01, *λ* = 0.05 (top). *M*-shape distribution of the *E* cells at stationary state for timescales ordering *τ*_*X*_ ≈*τ*_*N,−*_ *> τ*_*ξ*_ *> τ*_*N*,+_ with parameters *ϵ* = 0.5, *λ* = 1, *µ* = 0.01 (middle). Slow environmental switching regime, *τ*_*ξ*_ *> τ*_*X*_ *> τ*_*N*_, with parameters *ϵ* = 0.1, *λ* = 0.01, *µ* = 0.1 (bottom). Orange line in the tumor size distribution shows the diagonal terms in (5), see the text for discussion. **B)** Scattered plot of composition versus tumor size obtained from simulations for each row of A). The distribution of the tumor population size narrows with the increasing fraction of *E* cells in the tumor, see the discussion after (4).**C)** Difference of the timescales of tumor growth and composition change, Min[*τ*_*N ±*_] −*τ*_*X*_, for different values of symmetric phenotype switching and selection strength, *ϵ*. **D)** The support of the probability density function of the fraction of epithelial cells, (3), for different *µ* and *ϵ*.

In the slow-fluctuating regime, *τ*_*X*_ *< τ*_*ξ*_, there is sufficient time for the tumor composition to approach the equilibrium state in the current environment before the conditions switch. As a result, the stationary distribution accumulates near the boundaries, that is, equilibrium states 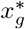 and 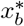, resulting in a bimodal distribution (see Fig.2A bottom-left panel). Indeed, for *τ*_*X*_ *< τ*_*ξ*_, the stationary probability density diverges at the endpoints, in contrast to the fast environment. In this slow switching regime, the stationary probability can assume an inverted *M*-shape as well, where *x*^*∗*^ becomes local maximum (see SI section IV for details). Nevertheless, the endpoints of the probability density of the tumor composition remain the most probable outcomes.

To obtain predictions for the tumor population size, we average the stationary pdf of the tumor population size for a fixed fraction of epithelial cells, (4), over all possible tumor compositions

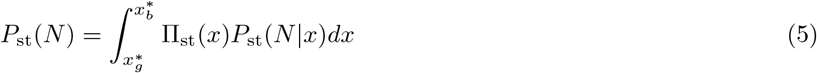

This construction provides a coarse-grained approximation of the total population size dynamics. Its validity, however, depends on the relative ordering of the characteristic timescales.

We first consider *ϵ <* 1 regime, whereby the variation of the reproduction rate of mesenchymal cells is smaller than its mean value.

When environmental fluctuations are faster than both tumor growth and phenotypic adaptation, *τ*_*ξ*_ *< τ*_*N*_, *τ*_*X*_, both stationary distributions (3) and (4) are typically unimodal. In the limit *τ*_*ξ*_ ≪ *τ*_*N*_, *τ*_*X*_, environmental fluctuations self-average and can be replaced by their mean value. For an unbiased environment this yields *N*^*∗*^ = *K* and *x*^*∗*^ = 1*/*2. The relative ordering of *τ*_*N*_ and *τ*_*X*_ determines the widths of stationary distributions, but their qualitative unimodal character is preserved.

The relative magnitude of *τ*_*X*_ and *τ*_*N*_ is controlled primarily by the phenotype switching rate *µ*. Increasing *µ* decreases the relaxation time of phenotypic adaptation, 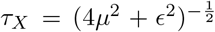, allowing the tumor composition to equilibrate faster than grows, for fixed *ϵ* (see Fig.2B). Meanwhile, the accessible range of tumor compositions narrows as a consequence of increased phenotype switching rates, since 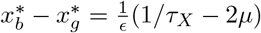, see Fig.2C. Thus, rapid phenotype switching reduces heterogeneity across tumors for a fixed selection strength *ϵ*.

When tumor growth is the fastest process, that is *τ*_*N*_ *± < τ*_*ξ*_ *< τ*_*X*_, the population size evolves as in the fixed environment and with fixed composition. The phenotypic adaptation, however, evolves under frequent switches of the environment since it is the slowest process in the considered case, resulting in a unimodal distribution of the composition with a most probable value *x*^*∗*^. While, the size distribution of the tumor is peaked around *K*(1 ± *ϵ*(1 −*x*^*∗*^)), as follows from (4) (see Fig.2A top-right panel).

For *ϵ* ∼ 1, the tumor size distribution can be unimodal even when environmental switching is no longer the fastest process (see Fig.2 middle-right panel). Here, the ordering of the characteristic timescales is *τ*_*N*+_ *< τ*_*ξ*_ *< τ*_*X*_ ≈ *τ*_*N,−*_ . In the distribution of the tumor population size, (4), the endpoint corresponding to *τ*_*N, −*_is suppressed for all values of the fraction of epithelial cells, *x*.

In the slow environment, *τ*_*X*_ *< τ*_*ξ*_, the distribution of the tumor composition peaks at the endpoints as discussed above. Nevertheless, the resulting distribution of the tumor size needs not to be bimodal.

The unimodal size distribution of the tumor can be observed when *τ*_*X*_ *< τ*_*ξ*_ *< τ*_*N*_ . The exponents in (4) are positive, ensuring that, for any given fraction of epithelial cells, the tumor size distribution is peaked inside the support. Here, the fraction of epithelial cells relaxes to its equilibrium value in the current environment before the environment switches, on average. Thus, from (3) and (4), it follows that the tumor population size (5) will be governed by (4) at the equilibrium state of the fraction of epithelial cells 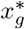 and 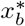, respectively. Therefore, satisfying the condition *τ*_*X*_ *< τ*_*N*_ results in reduced variability of tumors, again. Hence, the unimodality of the population size distribution is due to the rapid phenotypic adaptation resulting in the small support of the pdf of the tumor composition. At the crossover regime, *τ*_*N*_ ≈ *τ*_*ξ*_, the stationary distribution of the population size can become nearly uniform 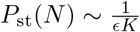 for small *ϵ* (see SI for further details).

When the environment switches slower than the tumor grows, *τ*_*ξ*_ *> τ*_*N*_, *P*_st_(*N\x*) and Π_st_(*x*) are bimodal because the exponents are negative in (4). However, simulations reveal two peaks for *P*_st_(*N*) rather than four as one might expect by analyzing *P*_st_(*N*| *x*) and Π_st_(*x*), see Fig.2A bottom-right panel. Indeed, for *τ*_*ξ*_ *> τ*_*X*_ *> τ*_*N*_, one may expect that the tumor size evolves toward its equilibrium value in the given environment, such that the composition of the tumor remains fixed during that time-interval. Thus, one can assume that for each possible value of *x*, there is always two peaks of (4) corresponding to *K*(1 *ϵ*(1 *x*)). Then, on intermediate time scales (*τ*_*X*_), the tumor composition evolves towards one of the possible values 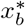 and 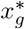, depending on the current state of the environment, thus, Π_st_(*x*) is bimodal as well. Hence, the expectation is that *P*_*st*_(*N*) might exhibits four peaks, that is *K*(1 *ϵ*(1 *x*_*i*_)), 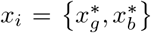 . The reason for the discrepancy between simulations and theory is that the peaks 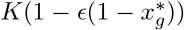 and 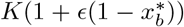 are not realized during the simulation, since this will correspond to the process where tumor size and composition evolve in effectively different environments, which is impossible.

The previous timescale arguments remain applicable for *ϵ >* 1 as well. However, in contrast to the *ϵ <* 1 case, here, *P*_st_(*N* |*x*) diverges at zero, that is, population extinction is possible, if 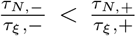(see SI section III for further details).

### Optimal phenotype switching rates in tumors

#### Optimal switching rates for long-term growth

The timescale analysis of (3) and (4) characterizes the evolutionary outcomes of the tumor growth and compositional changes. From the evolutionary perspective, it is important to understand out the role of phenotypic adaptation in a long-term growth of a tumor in stochastic environment [20]. In a resource-limited environment, the tumor size is always bounded, *N* ∈ 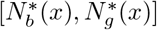 for all valid *x* although the possible size range depend on selection and phenotypic adaptation. We therefore focus on the mean population size of the tumor when the stationary state is reached.

A natural estimate of the mean tumor size is obtained directly from the stationary distributions (3)–(5). This approach predicts that the mean population size of the tumor is *K*(1 + *ϵδ*(1 − ⟨*x*⟩)), where ⟨*x*⟩ is the mean fraction of epithelial cells in the tumor which is calculated by averaging over its stationary distribution (3) (see SI section V for details). For unbiased environment, *δ* = 0, the mean population size of the tumor is predicted to be equal to the carrying capacity *K* of the environment. However, even for the unbiased environment, the simulation results disagree with the estimate unless the environment fluctuates sufficiently fast, *τ*_*X*_, *τ*_*N*_ *> τ*_*ξ*_, such that the environmental noise self-averages (see SI section V for further details).

A better estimate of the tumor size is obtained directly from the growth dynamics. According to (2), the instantaneous per-capita growth rate of the tumor is *R*(*x, ξ*) = 1 + *ϵξ*(1 − *x*). The per-capita growth rate is independent of the tumor size, hence it can be averaged exactly over the joint stationary distributions Π_st_(*x, ξ* = ±1), yielding the mean per-capita growth rate at the stationary state *E*_*X,ξ*_[*R*], that is, the long-term growth rate in the context of the model. Replacing the instantaneous growth rate with its mean value yields the effective logistic growth dynamics of the tumor with the effective carrying capacity *E*_*X,ξ*_[*R*]*K*, that is the upper bound for the average tumor size within the approximation. The average long-term growth rate, *E*_*x,ξ*_[*R*], is given by the following expression

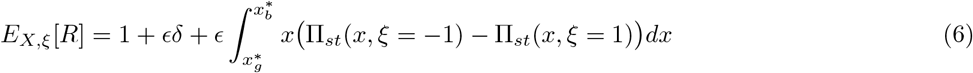

Even in unbiased environment, *δ* = 0, the long-term mean growth rate is not equal to the mean reproduction rate of the *M* and *E* cells (*E*_*ξ*_[*r*_*M*_] = *r*_*E*_ = 1 discussed above), in general, and increases with increasing *ϵ*. While decreases with increasing *µ* and *λ*, respectively (see SI section V for further analysis and derivation).

The mean long-term growth rate accurately reproduces the simulation results whenever environmental fluctuations are not too slow. In these regimes, the environmental state changes before the tumor reaches its equilibrium state in the current environment, such that the mean growth-rate reproduces the long-term dynamics of the tumor size. When the environment fluctuates rarely, that is *τ*_*ξ*_ ≫ *τ*_*X*_, then, both fraction of epithelial cells and the total population size converge to their stable equilibrium states in the current environment so that expressing the mean population size through the average long-term growth rate yields discrepancy between the simulation results and the estimate, *E*_*X,ξ*_[*R*]*K* (see SI section V). For large characteristic times of the environmental fluctuations the probability of observing tumor of size different than 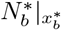 and 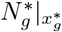 decreases and nullifies in the limiting case, corresponding to a quenched regime of the environment whereby the initial state of the environment governs the evolutionary outcome of the process. In this regime, the mean population size is 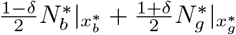

The optimal switching rates that provide the maximum long-term growth rate for the tumor population depend on *ϵ, δ* and *λ*. When *δ* = 0, the optimal phenotype switching rates decrease with decreasing *λ* and *ϵ*, see Fig.3A.

**FIG. 3:**
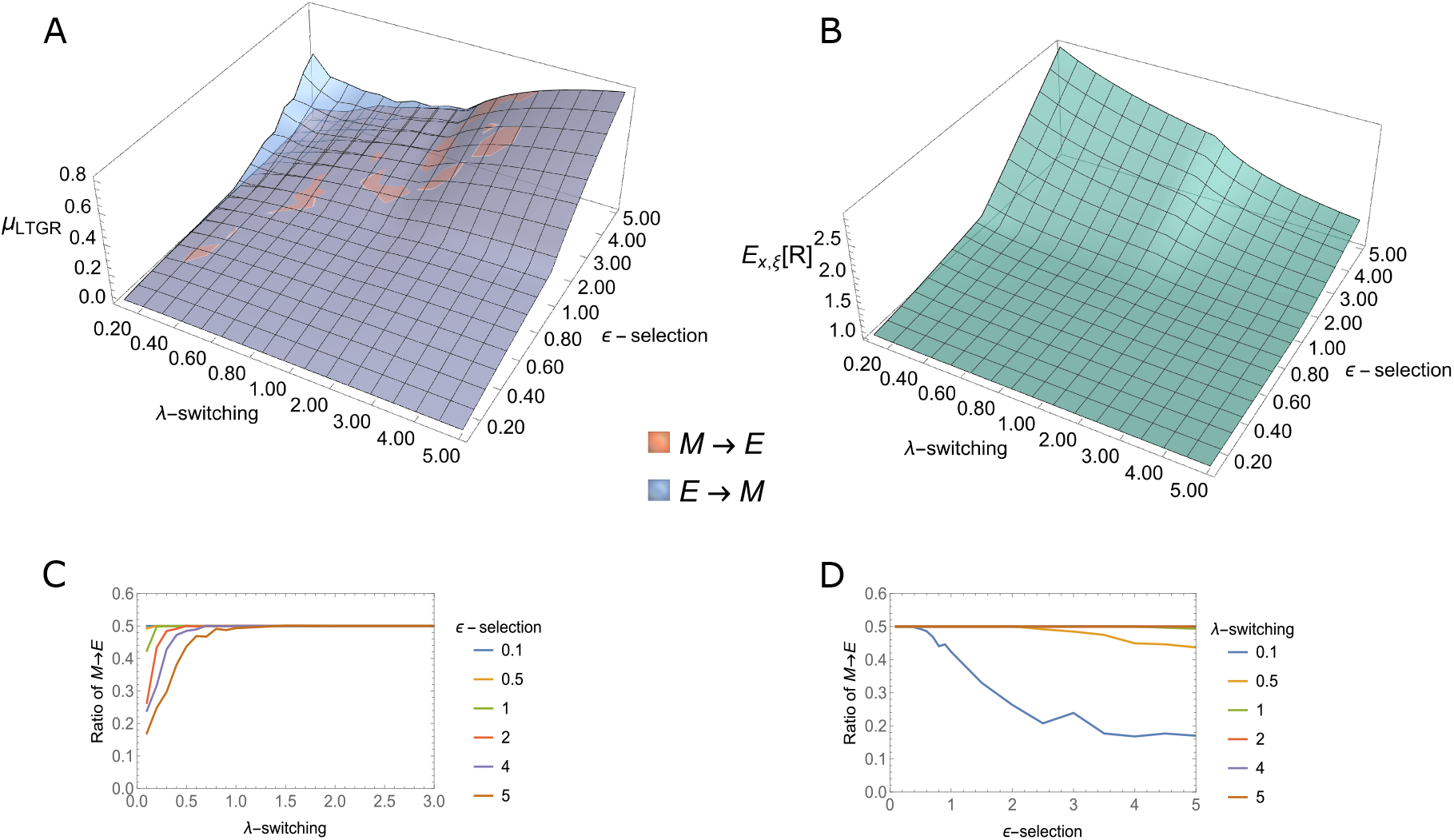
Optimal switching rates for long-term tumor growth rate for different model parameters, in an unbiased environment. **A)** Optimal switching rates for different environmental fluctuations rate, *λ*, and selective advantage and disadvantage of mesenchymal cells in favorable and unfavorable environment, *ϵ*,respectively. **B)** Long-term growth rate at optimal phenotype switching rates for each *ϵ* and *λ* at A). **C)** and **D)** Ratio of *M* → *E* switches in the tumor for various *ϵ* and *λ*.

For most combinations of the model parameters 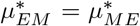, except in the slowly-switching environment with *ϵ >* 1, (Fig.3A). In this regime, the long-term growth rate of the tumor reaches its maximal value (Fig.3B). Here, phenotype switching from *E* → *M* is more favorable than *M* → *E* (Fig.3C and Fig.3D). Although the environment is unbiased, so that *M* cells have no selective advantage over *E* cells, on average, optimal switching rates allows the tumor to exploit the environmental fluctuations and grow. In the long periods of favorable environment tumor grows disproportionally due to the selective advantage of *M* cells and increased *M* → *E* switching compared to *E* → *M* . However, the optimal *E* → *M* switching rate is non-zero, because *M* cells alone will die out during the long periods of unfavorable environment. Indeed, neither *E* nor *M* cells alone are optimal for tumor proliferation in the stochastic environment. Thus, by tuning the phenotype switching rates, the tumor can exploit the advantages of both *E* and *M* phenotypes for growth, analogous to Parrondo’s paradox [46–48].

In a fast-fluctuating environment, the switching rates equalize,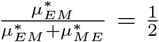 (Fig.3C). The long-term growth rate decreases as environmental fluctuations become more frequent, leading to self-averaging of the environmental noise, which nullifies in an unbiased environment, regardless of *ϵ*. Indeed, for greater values of *λ*, the environment self-averages, yielding a valley of possible switching rates, all resulting in the same long-term growth rate because the environment-dependent growth rate in (2) nullifies in the self-averaged regime. The balance between the phenotype switching rates is achieved easily in fast-fluctuating environments with *ϵ <* 1 (Fig.3C). However, in this regime, the optimal switching rates are lower than the switching rate of the environment, *λ*, indicating that optimal phenotype-switching does not require direct tracking of the environmental changes (Fig.3A).

Thus, in the considered scenario of temporally correlated environmental switches with resource-constrained tumor cell proliferation, the optimal switching rates do not tune to the statistics of environmental switches, *λ*, [4, 20, 21]. The non-monotonic behavior of the optimal switching rates is due to the non-linear dependence of the long-term growth rate, (6), on *ϵ* and *λ*, since in both the integration limits and the integrand depend on these parameters as well.

In the case of quenched environmental noise for the unbiased environment, *δ* = 0, whereby (6) does not accurately capture the tumor size, the average population size of the tumor is maximized when 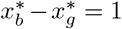 . The last condition holds, for any *ϵ*, when there is no switching between *E* and *M* cells at all, that is 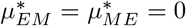 (see SI section V). Thus, on this regime, the switching between different cell types is deleterious.

The optimal switching rates define the characteristic timescale *τ*_*X*_ for a given *ϵ*. Nevertheless, phenotypic adaptation being neither faster nor slower than environmental fluctuations *τ*_*X*_ *> τ*_*ξ*_ and *τ*_*X*_ *< τ*_*ξ*_, respectively, is universal for optimal growth. That is, the ordering of the characteristic timescales at optimal switching rates depends on both *λ* and *ϵ*.

### Optimal switching rates for tumor survival

The above analysis describes the behavior of switching rates optimal for the long-term growth rate. Along with the long-term growth rate, another important target of tumor evolution is maximizing the survival time, that is, adapting to the stochasticity in the environment in such a way as to minimize the risk of the worst case outcome, tumor extinction due to large fluctuations. Therefore, long-term growth rate maximization does not necessarily reflect the safest evolutionary path for the tumor. Instead, the expected survival time can become the measure of tumor fitness.

We analyze the survival properties by considering the probability of reaching the minimal size of the tumor, that is the worst case outcome, compatible with survival and the expected time before this happens.

For a fixed composition of the tumor, *x*, the worst case outcome for the tumor is 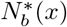, that is, the size of the tumor at equilibrium in the unfavorable environment with fixed fraction of *E* cells, (2). For 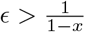, the worst case outcome is the extinction of the tumor, 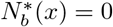 as follows from (2). Therefore, stochasticity of the environment and phenotypic adaptation of the tumor can prevent or delay the worst case outcome.

To quantify the risk of reaching the most unfavorable state, we introduce two artificial absorbing boundaries 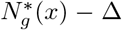 and 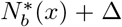, where Δ ≪ *K* is to ensure finite first-passage statistics. The risk minimization problem can therefore be formulated as a first-passage problem, that is studied for various dynamical systems under dichotomous noise ([49–51]).

The probability of hitting the boundary at left, that is 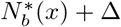, is denoted by *ψ*(*n, x*) (see SI section VI for the derivation). It depends on the initial population size, *n*, and the fraction of epithelial cells in the tumor, *x*. We first identify the composition of the tumor that minimizes the probability of the worst-case outcome, that 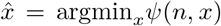. However, the fraction of epithelial cells *x* is not fixed and it evolves in the tumor due to phenotype switching and stochastic selection processes. We seek to identify the optimal phenotype switching rates such that the average of the fraction of epithelial cells, ⟨*x*⟩, is as close to 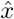 as possible in the given environment, linking the evolution of tumor composition to the risk minimization. Thus, 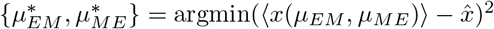. The resulting optimal switching rates are presented in Fig.4, where we assumed that initially the tumor population size is *K*. The dominant transition is *E* → *M* when *ϵ <* 1 (Fig.4A). At *ϵ* = 1, there is a sharp phase transition, where the *M* → *E* becomes dominant. The fraction of *M* → *E* transitions increases with increasing *ϵ*. This fraction is independent of the environmental fluctuation rate, *λ*, indicating that the magnitude rather than the frequency of the environmental fluctuations matters in this regime (Fig.4B,C). Thus, the risk of extinction is minimized by ‘escaping’ to the *E* state, which is less sensitive to environmental changes than the *M* state. Accordingly, MET is more important than EMT for avoiding tumor extinction in *ϵ >* 1.

**FIG. 4:**
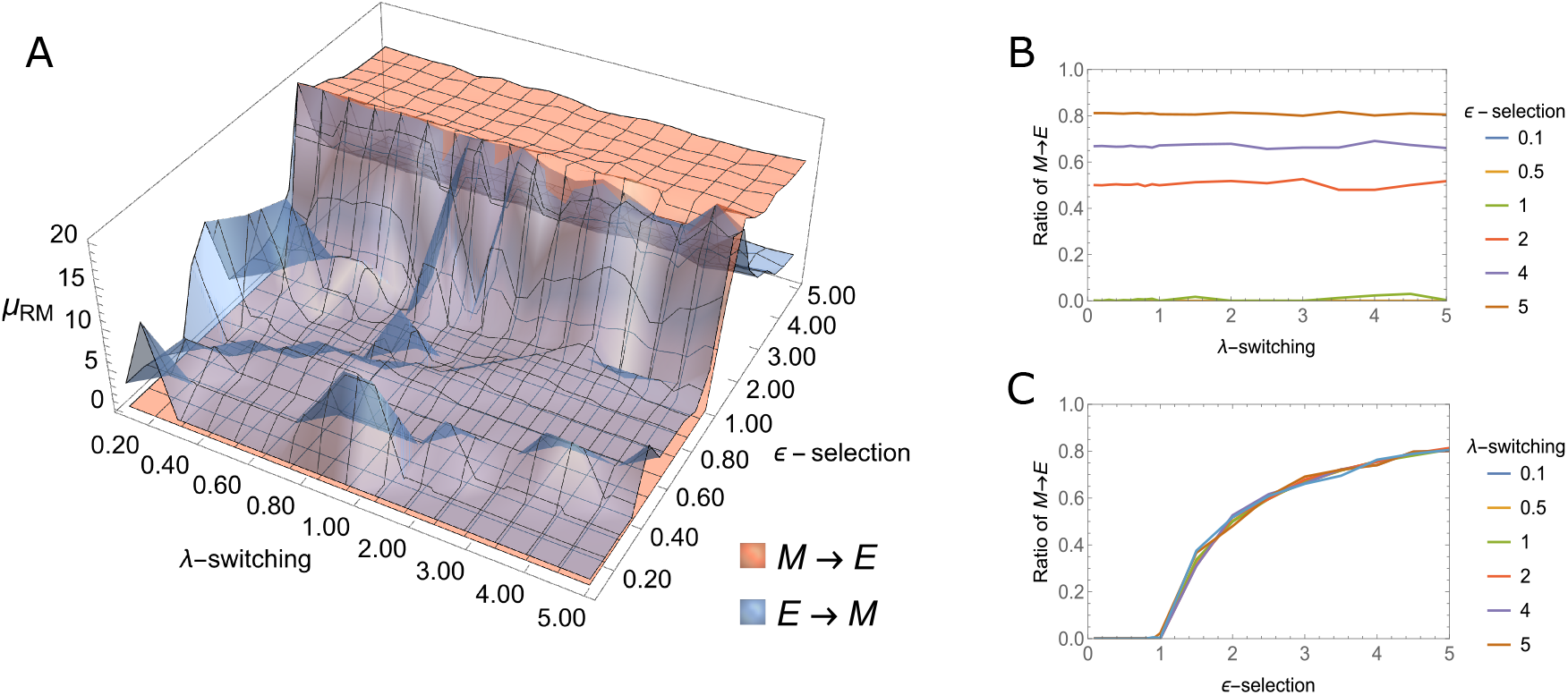
Optimal switching rates between epithelial and mesenchymal cells providing the minimal risk of tumor extinction and the maximal tumor survival time. **A)** The values of *E* →*M* and *M* → *E* switching rates for various environmental fluctuation rates, *λ*, and selection strength *ϵ*. The sharp phase transition occurs at *ϵ* = 1. **B** and **C)** The ratio of *M* → *E* transitions for optimal switching rates for different *ϵ* and *λ*.

Along with the risk minimization, we consider the maximum survival time of the tumor, *T* (*n, x*), assuming that 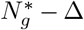 is a reflective boundary. Then, the maximal survival time for the tumor is the mean passage time to reach the only absorbing boundary at 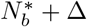 . The survival time of the tumor is also maximized for the same optimal composition 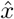 for the given model parameters. The tumor goes extinct in the bad environment in the case of *ϵ >* 1. Therefore, the extinction time diverges at 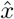 (see SI section VI).

### Implications for treatment

The maximum survival time of the tumor in an environment with *ϵ >* 1 is reached for the fraction of epithelial cells 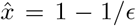 The adaptation of the tumor, then, involves tuning the phenotype switching rates such that the mean of the fraction of epithelial cells is the closest to the optimal fraction 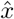 . These observations point to an optimal treatment strategy designed to drive the tumor composition away from the optimal state.

To this end, let us first consider tumor growth in fast-switching environment with *ϵ >* 1, that is *τ*_*N*_ *> τ*_*ξ*_, where the environmental noise self-averages. This regime is captured by (2) where the noise is substituted by the mean value, *ξ*(*t*) → *δ*.

In this regime, the tumor goes extinct whenever 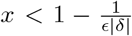, where *x* is the fraction of epithelial cells. Here, we consider the case of environmental bias towards the unfavorable state, *δ <* 0. If *δ >* 0, the tumor never goes extinct, as follows from (2) by substituting *ξ*(*t*) *δ* → *>* 0.

Therefore, to guarantee extinction for all admissible compositions of the tumor in the given environment, that is, for all 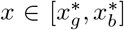, it suffices to impose extinction at the upper bound 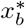. Specifically, if 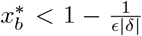 then, extinction would occur for the entire support 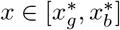

Solving this condition yields a threshold value for the MET switching rate, 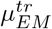, below which the tumor goes extinct

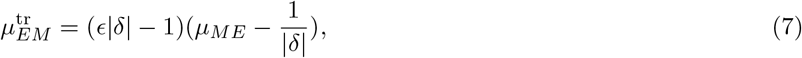

Note that (7) is valid for a harsh environment, such that *ϵ* | *δ*| *>* 1 yielding negative mean growth rate for *M* cells *E*_*ξ*_[*r*_*M*_] = 1 + *ϵδ <* 0.

The extinction of the tumor occurs whenever 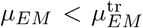 . Notably, the threshold increases linearly with the reverse switching rate *µ*_*ME*_, implying that, for given *ϵ* and *δ*, extinction of the tumor can be achieved either by suppressing *M* → *E* or increasing *E* → *M* transition such that 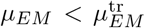 Moreover, increasing *ϵ* (treatment intensity), or environmental bias towards unfavorable state, |*δ*|, expands the extinction region, thereby reducing the required intervention on phenotype switching. The simulation results for time-evolution of the mean of tumor population size for fast environmental switching is shown in Fig.5A. In the simulations, we assume that extinction happens whenever the population size of the tumor is smaller than one. Tumor survives at phenotype-switching rates above the threshold value (7) (red line on Fig.5A). Increasing the phenotype switching rate, while not reaching the threshold value (7), results in the increase of the mean survival time of the tumor, with a wider range of trajectories (Fig.5A). The probability density function of extinction times of tumor is shown in the Fig.5B.

**FIG. 5:**
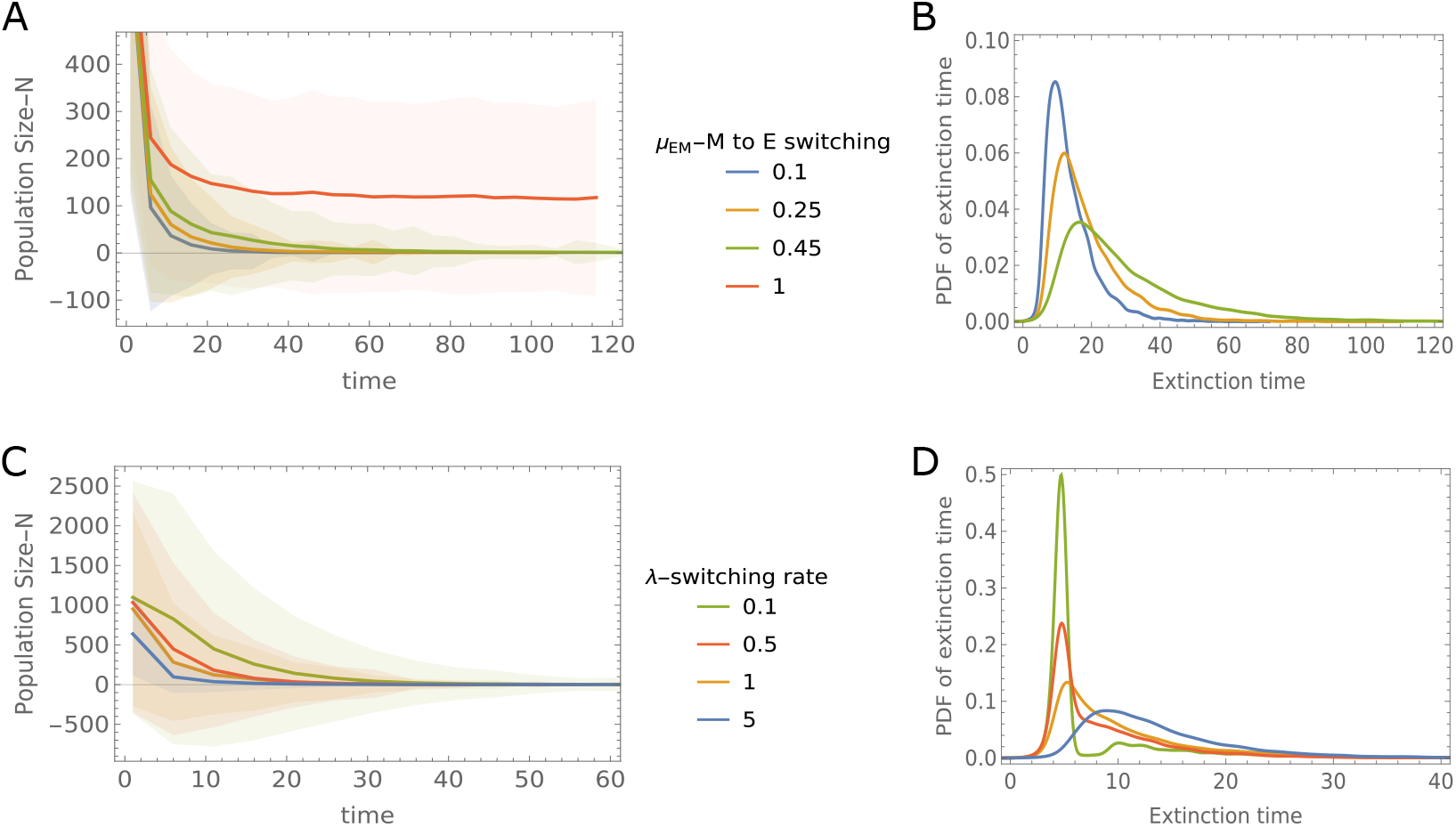
Tumor extinction. **A)** and **B)** Time evolution of the average population size of the tumor and probability density functions of extinction time, respectively, for different values of phenotype switching rate from *M* to *E* state, *µ*_*EM*_ in fast-fluctuating environment, *τ*_*ξ±,<*_ *τ*_*N, ±*_. The red line shows time-evolution of tumor size, for which 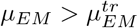.**C)** and **D)** Time evolution of tumor size and pdf of extinction time for various environmental switching rates. The phenotype switching rate from *M* to *E* state is below the threshold value. The model parameters are as follows, *ϵ* = *µ*_*ME*_ = 3, *δ* = −0.5, *K* = 1000, and the threshold value (7) is 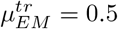. The results are obtained from 10^4^ independent runs. A), B) and C), D) are obtained for *λ* = 5 and *µ*_*EM*_ = 0.1 values, respectively.

The threshold value (7) is obtained under the assumption of a fast-fluctuating environment, but the obtained results hold for a slow-fluctuating environment, *τ*_*ξ*,_ *± > τ*_*N*,_ *±*, as well. Indeed, for *ϵ >* 1, the probability density function of the population size for the given fraction of epithelial cells, (4), is divergent near zero as long as 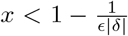for all environmental switching rates, *λ >* 0. Thus, the only evolutionary outcome in this case is the extinction of the tumor. The threshold value (7) is obtained for the greatest possible fraction of epithelial cells at any environment, 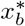, with no dependence on *λ*. Thus, the threshold is robust and applies for slow-switching environments as well. Nevertheless, we observe a qualitatively different behavior with increasing the environment switching rate *λ*.

In a slow-fluctuating environment, the mean population size decreases slower than in the fast-fluctuating regime, due to the rare long periods of favorable conditions, during which the tumor resumes growing again. Hence, the wider range of the time-evolution of the population size (Fig.5C). Increasing the switching rates of the environment smoothens the evolutionary trajectories of the population-size and the distribution of the extinction times, Fig.5D. In the case of slow environmental changes, the extinction times are concentrated at low values due to the likelihood of finding the tumor in the bad state at the initial stage, *δ <* 0. Thus, the tumor evolves towards extinction at longer times, but the environmental switches do not cause bursts of the tumor size.

## DISCUSSION

Motivated by experimental observations in cell lines, the impact of the variable microenvironment on tumor ecology [27, 52] and the paramount role of *E* − *M* phenotypic plasticity (EMP) in tumor evolution [33, 53], we developed a model of tumor proliferation in fluctuating environments, with stochastic phenotypic *E* − *M* switch. Despite its simplicity, our model captures rich, nontrivial behaviors of evolving tumors, with clinical, therapeutic implications (see Fig.6 for summary).

**FIG. 6:**
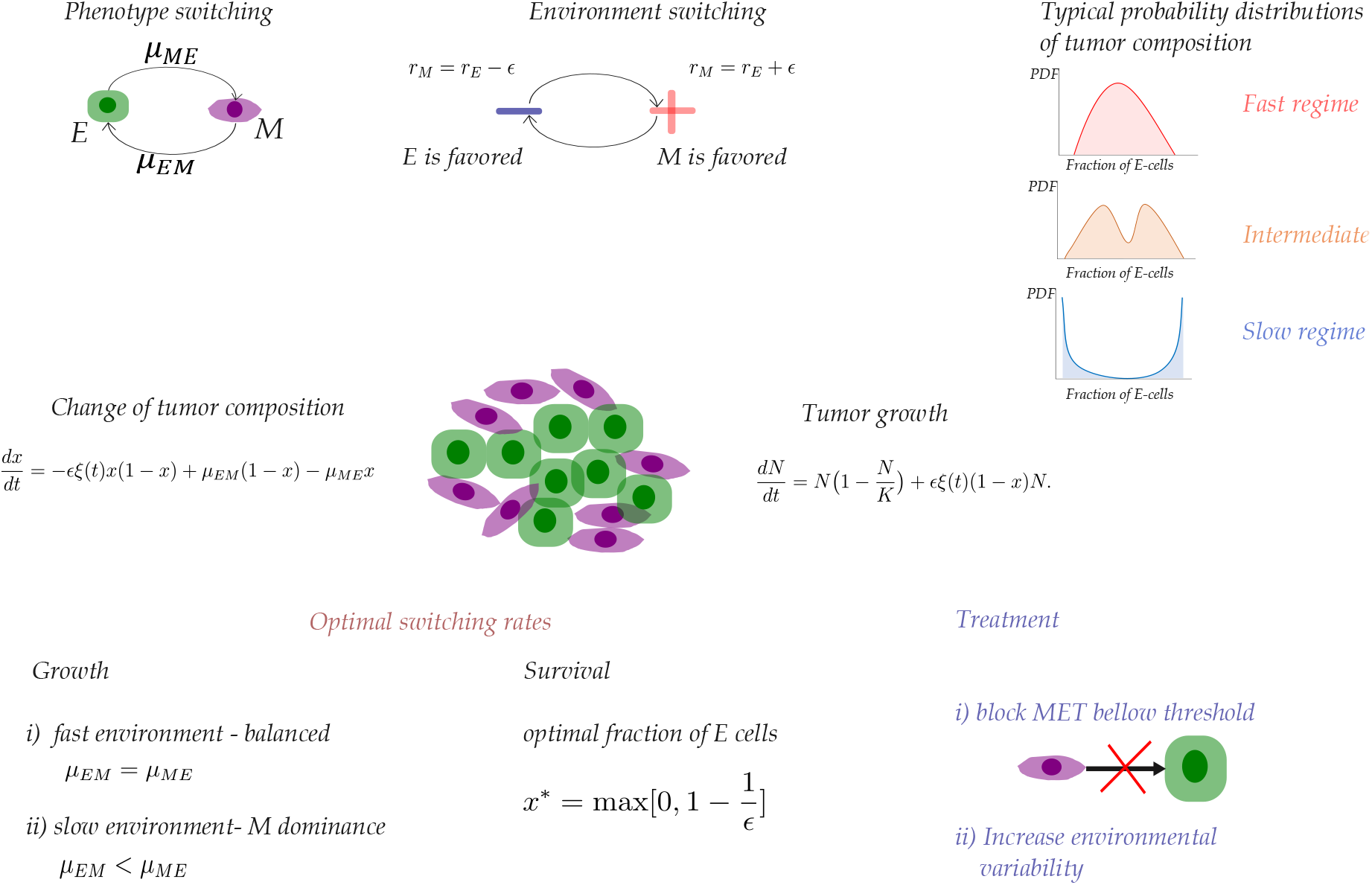
The role of EMT and MET in tumor growth and survival in stochastic environments: modeling framework and concepts. A phenotypically heterogeneous solid tumor, composed of epithelial (*E*) and mesenchymal (*M*) cells in a fluctuating environment, is depicted. Cells undergo reversible transitions between *E* and *M* states. Environment fluctuates between favorable and unfavorable states where *M* and *E* cells are advantageous, respectively. The evolutionary outcomes are defined by three characteristic timescales, corresponding to i) environmental fluctuations, ii) tumor growth (*N* (*t*) is the total number of cells in tumor) and iii) changes in the tumor composition (*x*(*t*) is the fraction of *E* cells in tumor). Changes in the relative ordering of these characteristic times result in various probability density functions for tumor composition and size: unimodal distribution in fast-fluctuating environments, multimodal distribution in intermediate cases and bimodal distribution in slow-fluctuating environments. The EMT/MET switching allows the tumor to optimize different fitness measures, namely, mean long-term growth rate or mean survival time of the tumor. The optimal phenotype switching rates differ for these fitness measures. In case of mean growth rate, the optimal switching rates are balanced in fast-fluctuating environment. However, EMT dominates over MET in slow-fluctuating environment with *ϵ >* 1, allowing the tumor to exploit fluctuating environment and growth. Conversely, for the maximal mean survival time, in *ϵ >* 1 environments, MET is increasingly favored over EMT in both fast and slow environment switching regimes with increasing *ϵ*. Optimal switching rate analysis yields insights into potential therapeutic interventions aiming at tumor extinction. Increasing the selective advantage of *M* cells over *E* cells in favorable environments, while increasing the vulnerability of *M* cells in unfavorable conditions and blocking MET below the threshold (7) leads to tumor extinction. In a favorable environment, *M* cells outcompete *E* cells due to their greater reproduction rate. However, blocking MET below threshold prevents the tumor (composed of *M*-cells dominantly) from escaping to the environment-independent *E* state. The threshold of MET depends on the environmental variability (via treatment frequency, intensity and scheduling) and EMT.

The evolutionary outcome of tumor proliferation and its phenotypic (i.e., *E* − *M*) composition is determined by three distinct processes and their relative timescales, namely, environmental changes, phenotypic adaptation via stochastic switching and selection, and population growth. The interplay between processes leading to a rich phase plane structure. The analysis of the model reveals coupling between phenotypic composition and tumor size, whereby tumor size is governed by the phenotypic composition, but phenotypic heterogeneity is independent of the tumor size. In a fast-fluctuating environment, both phenotypic composition and tumor size evolve slower compared to environmental changes, such that the averaged environmental conditions determine the evolutionary outcomes, with the tumor phenotypic composition and population size exhibiting unimodal distributions around the deterministic equilibrium state. Conversely, in a slow-fluctuating environment, tumor composition evolves towards equilibrium in the current environment and concentrates around the equilibrium values of the quenched disorder limit of the environment, resulting in a bimodal distribution of phenotypic composition. Between these extremes, intermediate regimes generate nontrivial distributions of tumor composition that cannot be inferred from fitness landscapes of fixed environments alone, such as M-shape and inverted M-shape distributions in fast and slow environments, respectively, whereby the extrema of the distributions are not linked to the proliferation of the tumor in a fixed environment. Further, we obtained the phase plane for the population size distribution using the adiabatic approximation [37]. The ordering of the characteristic timescales of tumor growth and composition defines the range of phenotypic heterogeneity of the tumor in a stochastic environment. In particular, the extent of tumor heterogeneity increases and decreases when the tumor growth is slower and faster than phenotypic adaptation, respectively (Fig.6).

From an evolutionary perspective, the analysis of our modeal reveals a fundamental trade-off between growth optimization and extinction risk minimization. Maximizing the long-term growth rate of the population in a stochastic environment captures the long-term evolutionary success of the population in resource-unconstrained models [4, 20, 21]. Here, we first identify the long-term growth rate of the population in a resource-constrained environment. Optimizing this quantity over phenotype switching rates reveals that, in a fast-fluctuating environment, the optimal switching rates are balanced when the environment is unbiased. In this case, the optimal switching strategy becomes independent of the current environmental state, so that it is equally likely to observe EMT and MTE switching, irrespective of the selection pressure of the environment, revealing the bet-hedging strategy of the tumor cell population. Moreover, the switching rates depend on the selection pressure acting on the cells in the current environment, *ϵ*: increasing *ϵ* increases the phenotypic switching rates such that the population neither suffers nor benefits from environmental switches. In a slow-fluctuating environment, the symmetry between the optimal switching rates breaks, favoring EMT, that is, switching toward phenotypes that are environment-sensitive (i.e., *M* cells), thereby maximizing the long-term growth rate. Therefore, in a quenched environment, where the stochasticity of the environment enters only due to randomly chosen initial state, phenotype switching becomes deleterious for the long-term growth of the tumor (Fig.6).

In contrast, minimizing the extinction risk, and equivalently maximizing the survival time of the tumor, leads to qualitatively different optimal phenotype switching rates. Both quantities attain their optima at the same tumor composition. This optimal composition corresponds to a purely mesenchymal population in *ϵ <* 1 environments. In the opposite case, *ϵ >* 1, the optimal composition is such that the tumor size remains constant during unfavorable periods (owing to the insensitivity *E* cells to the environment), whereas during favorable periods, the tumor grows following environmental switches (owing to the sensitivity of *M* cells to the environment). The optimal phenotype switching rates are those that keep the average tumor composition as close as possible to this optimal composition. The fraction of MET in the population increases with increasing *ϵ* (Fig.6).

The framework developed here has several nontrivial implications for potential cancer treatment. Specifically, we observe a sharp phase transition in the optimal switching rates at the crossover from mild environments (*ϵ <* 1) to harsh environments (*ϵ >* 1). Tumor extinction is possible only in *ϵ >* 1 environments, implying that, for tumor extinction, large fluctuations in the selective advantage of *M* cells are necessary. In harsh environments, tumor survival critically depends on the *E* cells because they are insensitive to the environment, whereas the *M* cells are vulnerable. Thus, blocking MET becomes crucial, as it prevents te tumor from escaping to the environmental-insensitive *E* state. In particular, in harsh environments, the MET to EMT ratio increases with increasing fluctuations in reproduction rates, thereby maintaining the optimal phenotypic composition in the given stochastic environment for the survival of the tumor. Indeed, we derive the threshold value of MET switching rate below which the tumor cannot survive in a harsh environment that is biased toward a state favoring *E* cells (7). This *M* → *E* switching rate threshold is a target for intervention. The threshold value links the EMT rate, fluctuations in the reproduction rate of the *M* cells that are controlled by the treatment intensity, and the environmental bias-dosage scheduling. Notably, this threshold is independent of the environmental switching rate, ensuring tumor extinction in both slow and fast environments. Therefore, treatment strategies aiming to inhibit MET can remain effective even under poorly controlled or unpredictable conditions, where the environmental timescales cannot be accurately controlled. However, the extinction-path dynamics differ between environments; in a slow environment, a burst in tumor size might occur because of rare sequences of environmental states favoring mesenchymal cells. In contrast, in the fast-fluctuating regime, such bursts are not observed, and the tumor goes extinct without abrupt changes in size. Thus, blocking MET coupled with increased environmental variability (via treatment frequency, intensity and scheduling) is expected to be the most efficient therapeutic intervention. This approach can be realized by targeting transcription factors that play pivotal roles in EMP, such as GRHL2, which epigenetically regulate the dynamic extent of *E* and *M* states [38, 54, 55].

The broader implication of this study is that tumor evolution in fluctuating environments cannot be understood without explicitly accounting for dynamical coupling between tumor phenotypic composition and tumor size, and the timescales that govern these processes. This study shows that even a low-dimensional stochastic model of tumor evolution can generate rich and clinically relevant behavior when the details of proliferation processes in the stochastic environment are explicitly incorporated into the model. Nonetheless, further development of the model might provide additional insights into tumor evolution and how it can be exploited for therapeutics. The model can be expanded to several directions, in particular, addressing the capacity of cells to migrate and invade. Such expansion potentially could further elucidate the role of EMP in tumor progression to distant metastatic regions and suggest improved therapeutic strategies [56]. The current model does not take into an account the demographic noise in the tumor proliferation, which can affect the fixation of phenotypes within the tumor as well as tumor extinction. Furthermore, in the current model, the environment is represented by stationary noise, with time-independent switching rates. However, the tumor itself can alter it’s own microenvironment, and thus, the environmental noise statistics as well. Further understanding how the variable environment can affect tumor initiation and progression would require integrating the complex mutational processes into tumor evolution models, aiming to set a theoretical framework to investigate the rich interplay between environmental changes, genetics and non-genetic mechanisms [57, 58].

## METHODS

### Generation of MCF10A-derived tumorigenic clones

To model cellular adaptation to fluctuating nutrient availability, MCF10A non-tumorigenic mammary epithelial (*E*) cells were subjected to long-term selection under cyclical growth and nutrient depletion conditions, whereby these cells acquire driver mutation and become neoplastic, as previously described [38]. Cells were cultured in T75 flasks with 15 mL of DMEM/F12 medium supplemented with EGF (10 ng/mL), hydrocortisone (0.5 µg/mL), cholera toxin (100 ng/mL), insulin (5 µg/mL), 5% horse serum, and 1% penicillin–streptomycin. Briefly, cells were cultured without media replacement until culture confluency declined to approximately 50% as a consequence of progressive nutrient and cytokine depletion, media acidification and confluency, leading to spontaneous cell death. At this point, culture media was replaced and surviving cells were allowed to regrow. This process was repeated continuously for approximately two years. At the end of the selection period, clones were isolated by limiting dilution and expanded. Based on their morphology as well as their transcriptomic and epigenetic states, the tumorigenic clones were classified as either *E*-dominant (PE14 and PE16) or mesenchymal (*M*)-dominant (PE21 and PE26) [38].

### Longitudinal imaging of tumorigenic clones during nutrient depletion

Parental MCF10A cells and 4 tumorigenic clones (PE14, PE16, PE21, PE26) were seeded into T75 culture flasks at a density of 110^6^ cells per flask in 15 mL of complete growth medium. Following cell attachment, cultures typically exhibited approximately 25% confluency. Cells were maintained under standard incubation conditions (37°C, 5% *CO*_2_). Beginning on the day of seeding, cultures were monitored longitudinally by brightfield phase contrast microscopy using an EVOS FL Auto 2 imaging system (Thermo Fisher Scientific) equipped with a 4*X* objective. For each flask, a 4 × 4 matrix of images (16 fields total) was acquired at five focal planes of the flask every other day to capture representative population-level morphology and confluency. The culture medium was intentionally not replaced in this phase of the study, allowing nutrients to become progressively depleted and metabolic waste products to accumulate over time. Cells were imaged until confluency declined to approximately 50 %, as determined by visual inspection. At this point, the medium was replaced with 15 mL of fresh growth medium, allowing the surviving cells to recover and re-expand, thereby initiating a new selection cycle. Representative images corresponding to the initial seeding stage, full confluency, onset of confluency loss, and approximately 50 % confluency were selected for downstream analysis. Temporal confluency measurements were used to characterize clone-specific responses to progressive nutrient depletion. For reference, the same experiment was conducted for each of the clones with media being replenished every two days.

### Image-based quantification of viability of MCF10A Parental and tumorigenic clones

Longitudinal brightfield images were analyzed for each replicate (*n* = 16) at five focal planes, each 8*m* apart (z00, z01, z02, z03, z04) of the flask for each time point in MATLAB to quantify time-dependent changes in areas occupied by live cells across MCF10A parental and tumorigenic clones. Images from each replicate were read with imread function, converted to grayscale with rgb2gray function, and converted to double precision with im2double function. Sequential frames were aligned by phase-correlation-based image registration using fft2, ifft2, and imtranslate, followed by cropping to the common valid image region. For each frame, uneven illumination and background structure were corrected using imflatfield, mat2gray, and imgaussfilt. Live cell-associated foreground regions were identified using combined scores of changes in intensity (contrast), texture (standard deviation of neighboring pixels), and edge features (gradient estimation around a pixel – edges have sharp gradients) generated with imbothat, imtophat, imgradient, and stdfilt, followed by morphological refinement using strel, imclose, imdilate, bwareaopen, and regionfill. This refined image is used to identify connected components using bwconncomp function, identifying distinct objects that have similar intensity pixels clustered together as a connected component. These connected components are classified into two categories, cluster of adherent cells and isolated individual cells, based on the proximity between connected components (SI section I Supplementary Figure 1). The final live-cell area was calculated as the percent of the analyzed image region occupied by live cell-associated foreground (SI section I Supplementary Figure 1). This was repeated for 16 replicates for each z-position and quantified as mean ± standard deviation over time. Time was expressed as days elapsed from the earliest imaging date in each experiment. The transition from nutrient deprivation to fresh media was marked using a vertical dashed line (Fig.1C-D). This analysis was applied to parental MCF10A cells and tumorigenic clones imaged longitudinally under media-changing and starvation/re-feeding conditions.

## AUTHORS CONTRIBUTIONS

S.G.B, Y.I.Y. A.S.S., E.V.K and E.P. conceived and designed the study. P.R.S., R.R.C., A.S.S. designed, performed and analyzed the experimental work. S.B. developed and performed the analytical and computational theoretical work. All authors analyzed and interpreted the data. S.B., E.V.K. and E.P. wrote the manuscript, with contributions from all authors. All authors have read and approved the final version of the manuscript.

## ACKNOWLEDGMENTS

We are grateful to Vahagn Abgaryan, Vardan Bardakchyan and Koonin group members for useful discussions. S.G.B is supported by Higher Education and Science Committee of RA (Research Program 24IRF/2-1C001). The experimental work was primarily funded by the Physical Sciences–Oncology Network (PS-ON) grant U01CA261841. Moffitt Core facilities were supported by the Cancer Center Support Grant P30-CA076292. Time-lapse imaging of the MCF10A and PE clones was performed using an EVOS FL Auto 2 system, acquired through philanthropic support from the Pentecost Family Myeloma Research Center, to which the authors express their sincere gratitude. E.P., Y. I.W and E.V.K. are supported by the Intramural Research Program of the National Institutes of Health (NIH). The contributions of the NIH author(s) are considered Works of the United States Government. The findings and conclusions presented in this paper are those of the author(s) and do not necessarily reflect the views of the NIH or the U.S. Department of Health and Human Services.

