## Supplementary Information for "Evolutionary implications of phenotype switching for growth and survival of tumors in stochastic environments"

### I. SUPPLEMENTARY FIGURES

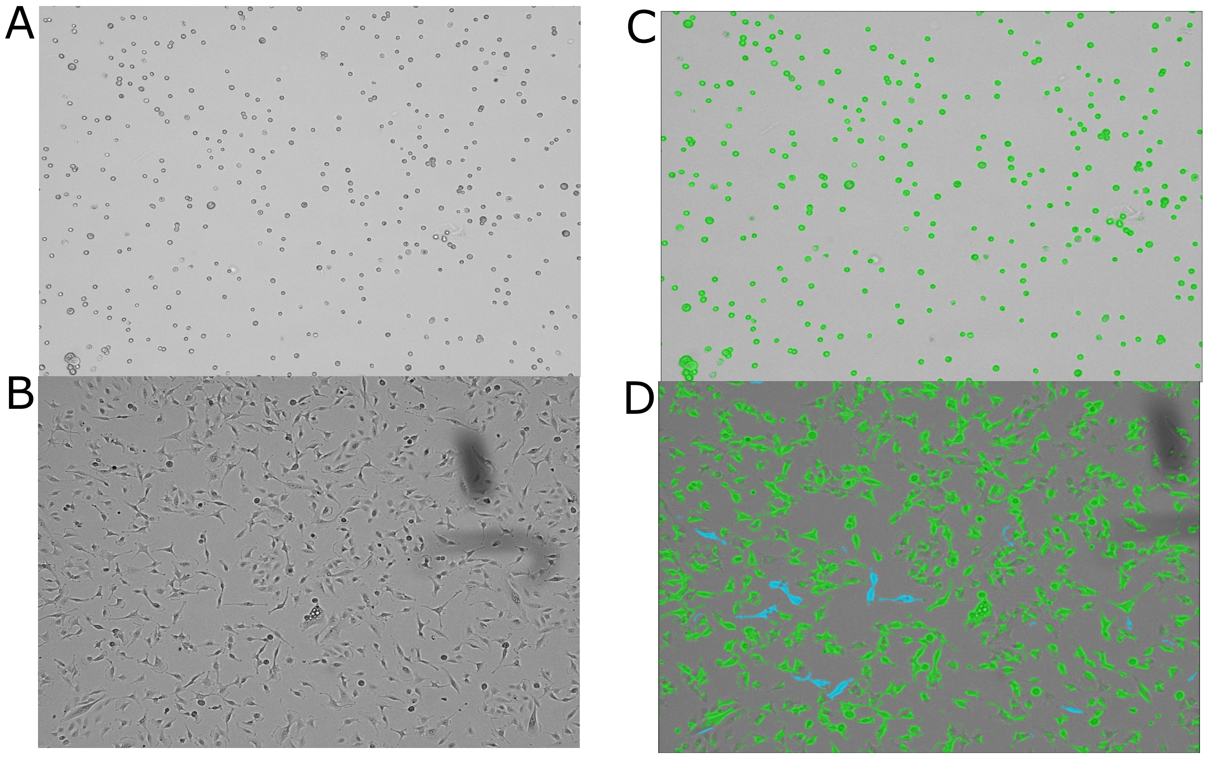


Supplementary Figure 1: **Brightfield images and overlays of live cell areas.**
(A) and (B) show brightfield images at initial seeding and increased confluence at the next time point. (C) and (D) show overlays of live cell (green/cyan) areas identified by the algorithm described in Methods. Green overlay indicates adherent cells and cyan overlay indicates islolated individual cells..

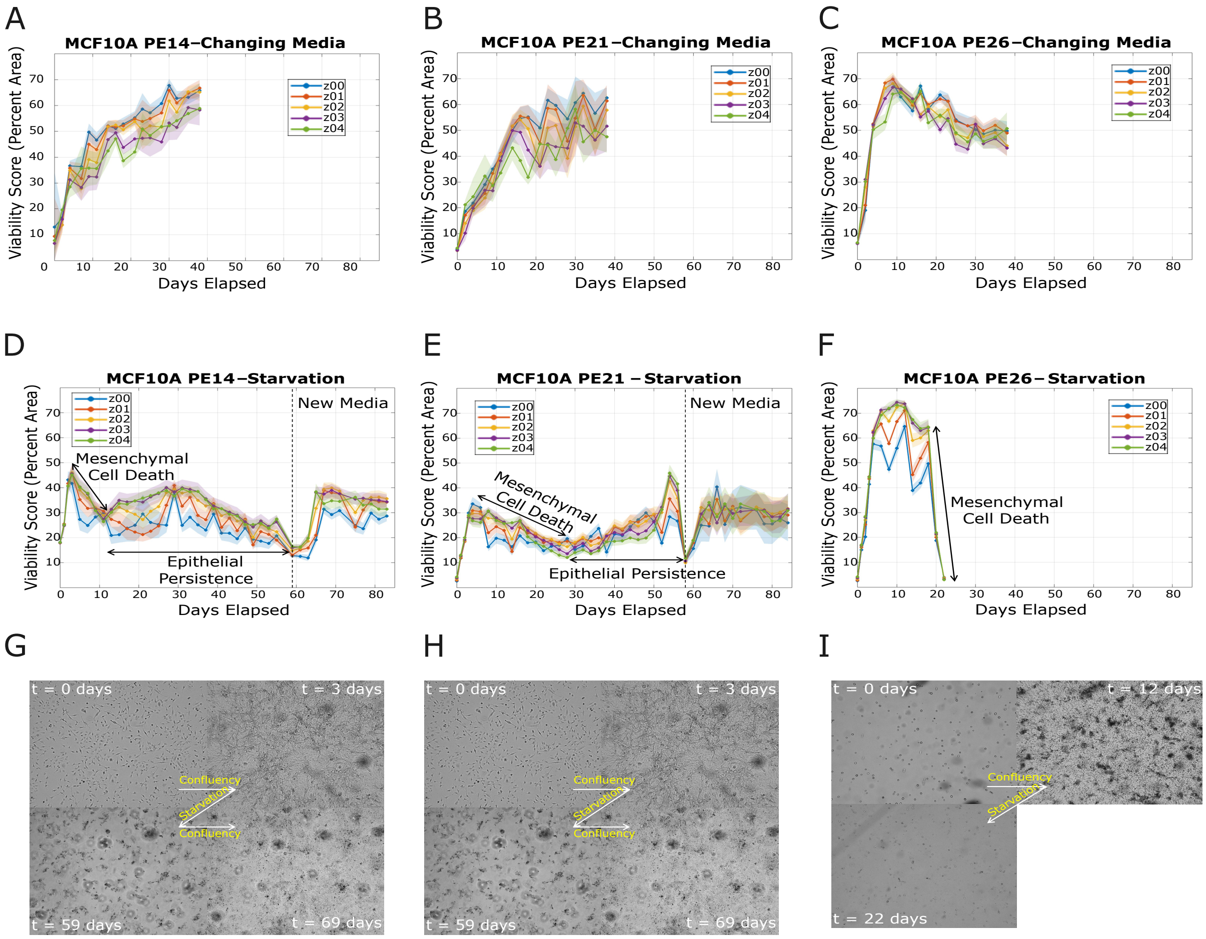


Supplementary Figure 2: **Nutrient deprivation induces a biphasic response in tumorigenic clones PE14, PE21, and PE26.**

(A), (B), and (C) show cells maintained under media-changing conditions for PE14, PE21, and PE26 clones, respectively. Under these nutrient-replete conditions, areas occupied by live cells generally increased or remained stable, reflecting continued growth and maintenance of adherent cultures. (D), (E), and (F) show matched starvation/re-feeding conditions. (G), (H), and (I) show representative brightfield images illustrating the transition from initial seeding to confluency, starvation-associated loss of live cells, persistence of a subpopulation, and subsequent confluency upon media replenishment corresponding to the experimental conditions D, E, and F, respectively. We note that in (C) the M cells can grow on top of the existing E/M cells, forming dense 3D structures; which cannot be accurately quantified in a 2D imaging system (Methods). The apparent decline in measured viability in (C) may be due to the transformation of a planar structure (x-y axes) to a 3D structure (x-y-z axes), while the M cells continue to proliferate in this nutrient-rich condition (i.e., frequent replenishment of media).

### II. POPULATION DYNAMICS. CHARACTERISTIC TIME-SCALES.

In the main text we describe the time evolution of the tumor through the fraction of epithelial cells, $x$, and the tumor population size, $N$.

The equations are derived from the following dynamical system

$$\begin{aligned} \frac{dn_E}{dt} &= n_E(r_E - \frac{n_E + n_M}{K}) + \mu_{EM}n_M - \mu_{ME}n_E, \\ \frac{dn_M}{dt} &= n_M(r_M - \frac{n_E + n_M}{K}) - \mu_{EM}n_M + \mu_{ME}n_E, \end{aligned} \tag{1}$$

where $n_E$ and $n_M$ are the population size of $E$ and $M$ cells in the tumor, respectively. Introducing the variable fraction of epithelial cells, $x = \frac{n_E}{n_E+n_M}$, and total population size, $N = n_E + n_M$, and using (1) we get

$$\frac{dx}{dt} = x(1-x)\big(r_E - r_M\big) + \mu_{E\leftarrow M}(1-x) - \mu_{M\leftarrow E}x, \tag{2}$$

$$\frac{dN}{dt} = N\big(r_E x + r_M(1-x) - \frac{N}{K}\big), \tag{3}$$

In the absence of a phenotype switching between different type of cells, that is, $\mu_{EM} = \mu_{ME} = 0$, the tumor will be homogeneous in a stable steady state, with only epithelial or mesenchymal cells. The outcome depends on the difference in the reproduction rates $r_E - r_M$, when $r_E = r_M$ the composition of the tumor does not evolve over time.

If the change of tumor composition is governed by the phenotype switching process only, that is $r_E = r_M$, then the fraction of epithelial cells at the steady state is $x^* = \frac{\mu_{EM}}{\mu_{EM}+\mu_{ME}}$, while the total population size of the tumor is $N^* = r_E K$.

In the more general case, that is $r_E \neq r_M$ and $\mu_{EM}, \mu_{ME} \neq 0$, the dynamical system (2,3) represents the interplay between selection and phenotype switching mechanisms that together govern the time evolution of tumor. In the presence of environmental fluctuations the composition and tumor size evolve as follows

$$\frac{dx}{dt} = -\epsilon\xi(t)x(1-x) + \mu_{em}(1-x) - \mu_{me}x, \tag{4}$$

$$\frac{dN}{dt} = N\big(1 - \frac{N}{K}\big) + \epsilon\xi(t)(1-x)N. \tag{5}$$

**II.A. Characteristic times.**

To figure out the characteristic timescales of the size and the composition changes of the tumor we analyze (2) and (3) in fixed environment $\xi = \pm 1$.

In both good and bad environment the right-hand side of (2) can be represented in the following form

$$\frac{dx}{dt} = ax^2 + bx + c \quad (6)$$

where $a = \pm\epsilon$, $c = \mu_{EM}$ and $b = -(\mu_{EM} + \mu_{ME} \pm \epsilon)$ for good and bad environment $\xi = \pm 1$.

Using above introduced notations we transform the right hand-side of (6) into $a(x-x_1)(x-x_2)$, where $x_{1,2}$ are the solutions of $ax^2 + bx + c = 0$ in the given environment, that is $x_{1,2} = \frac{-b \pm \sqrt{b^2-4ac}}{2a}$.

From (6) we obtain

$$\int \frac{dx}{a(x-x_1)(x-x_2)} = t + C \quad (7)$$

The integrand in (7) is equal to $\frac{1}{a(x-x_1)(x-x_2)} = \frac{1}{a(x_1-x_2)}(\frac{1}{x-x_1} - \frac{1}{x-x_2})$. Plugging back the last relation into (7) we obtain that

$$\ln|\frac{x-x_1}{x-x_2}| = a(x_1-x_2)t + C = -\sqrt{b^2-4ac}\, t + C \quad (8)$$

Thus, the time evolution of the fraction of $E$ cells in the tumor converges exponentially to its steady state with a rate defined by the discriminant of the right hand-side of (6). Then, the characteristic timescale of $x(t)$ is $\tau_X \propto 1/\sqrt{b^2-4ac}$.

The time-evolution of tumor population size is described by the logistic growth model with modified reproduction rate, that depends on the fraction of epithelial cells in the tumor, that is

$$\frac{dN}{dt} = N(\alpha - \frac{N}{K}), \quad (9)$$

where $\alpha \equiv 1 + \epsilon\xi(1-x)$. We assume that the fraction of epithelial cells is fixed, then the total population size converges to its steady state in the given environment, that is $\max[K(1-\epsilon(1-x)), 0]$ and $K(1+\epsilon(1-x))$ in bad and good environment, respectively. The convergence is again exponential with a rate $|\epsilon(1-x)-1|$ and $1+\epsilon(1-x)$, hence the characteristic timescale of the time evolution of population size is $\tau_{N,\pm} \propto \frac{1}{|1\pm\epsilon|}$.

### III. STATIONARY DISTRIBUTIONS FOR TUMOR SIZE AND COMPOSITION

Here, we provide details on the derivation of stationary probability functions for the composition and size of the tumor. To this end let us consider an auxiliary variable $z$, such that the time-evolution of it is governed by following partially-deterministic Markov Process

$$\frac{dz}{dt} = f(z) + \xi(t)g(z) \tag{10}$$

Here it is assumed that in each environment there is only one stable state, that solves the following equations $f(z^*) \pm g(z^*) = 0$.

The stationary probability density functions are found by solving forward-Kolmogorov equations for the joint probability functions, which are denoted by $\rho(z, t, \xi = \pm 1)$. The forward-Kolmogorov equations for the joint probability functions are as follows (see [1] for derivation)

$$\partial_t \rho(z, t, \xi = 1) = -\partial_z \Big( \big(f(z) + g(z)\big)\rho(z, t, \xi = 1) \Big) + \lambda^- \rho(z, t, \xi = -1) - \lambda^+ \rho(z, t, \xi = 1), \tag{11}$$

$$\partial_t \rho(z, t, \xi = -1) = -\partial_z \Big( \big(f(z) - g(z)\big)\rho(z, t, \xi = -1) \Big) - \lambda^- \rho(z, t, \xi = -1) + \lambda^+ \rho(z, t, \xi = 1), \tag{12}$$

where $\lambda^{\pm} = \lambda(1 \mp \delta)$ is the switching rate from good to bad states and vice versa. In the stationary state, that is $\partial_t \rho_{st}(z, \xi = \pm 1) = 0$, summing up the right hand sides of (11) and (12), and get the following relation between the stationary distributions

$$\big(f(z) + g(z)\big)\rho_{st}(z, \xi = 1) + \big(f(z) - g(z)\big)\rho_{st}(z, \xi = -1) = \text{const} \tag{13}$$

The constant in (13) is set to zero due to natural boundary conditions on the probability density functions on the infinity.

From (11) and (12) for the stationary distributions we obtain the following expressions

$$\rho(z, \xi = \pm 1) = \frac{1}{Z} \frac{1}{|f(z) \pm g(z)|} e^{\int_u^z \frac{\lambda^-}{f(y) - g(y)} + \frac{\lambda^+}{f(y) + g(y)} dy} \tag{14}$$

in the derivation we used the relation (13) between the stationary probability densities, $u \in [z_-^*, z_+^*]$ is arbitrarily. The constant $Z$ is for normalization of the joint probability densities, that is found from the following expression $\frac{1}{Z} \int_{z_-^*}^{z_+^*} \rho(z, \xi = 1) + \rho(z, \xi = -1) dz = 1$.

For the tumor population size $f(z) = z(1 - \frac{z}{K})$ and $g(z) = \epsilon(1-x)z$ are inserted in (10), while for the fractions of epithelial cells in the tumor $f(z) = \mu_{EM}(1-z) - \mu_{ME}z$ and $g(z) = -\epsilon z(1-z)$, and after integrations obtained the expressions

$$\Pi_{\rm st}(x) = \Pi_{\rm st}(x, \xi = -1) + \Pi_{\rm st}(x, \xi = 1) = \\ = \frac{1}{Z_x}\left(\frac{1}{(x_b^* - x)(x - x_{b,2}^*)} + \frac{1}{(x - x_g^*)(x_{g,2}^* - x)}\right)\left(\frac{x_b^* - x}{x - x_{b,2}^*}\right)^{\frac{\tau_{x,-}}{\tau_{\xi,-}}}\left(\frac{x - x_g^*}{x_{g,2}^* - x}\right)^{\frac{\tau_{x,+}}{\tau_{\xi,+}}}, \quad (15)$$

$$P_{\rm st}(N|x) = \frac{1}{Z_N(x)}\frac{1}{N^3}\left(\frac{K(1+\epsilon(1-x)) - N}{N}\right)^{\frac{\tau_{N,+}}{\tau_{\xi,+}}-1}\left(\frac{N - K(1-\epsilon(1-x))}{N}\right)^{\frac{\tau_{N,-}}{\tau_{\xi,-}}-1} \quad (16)$$

The normalization constant $Z_N(x)$ is found by integrating the distribution function over the support $[K(1-\epsilon(1-x)), K(1+\epsilon(1-x))]$, where we assumed $\epsilon < 1$.

$$Z_N(x) = \int_B^A \frac{1}{N^3}\left(\frac{A-N}{N}\right)^{\alpha-1}\left(\frac{N-B}{N}\right)^{\beta-1} dN = \frac{A^{-1-\beta}B^{-1-\alpha}(A-B)^{-1+\alpha+\beta}(A\alpha + B\beta)\Gamma[\alpha]\Gamma[\beta]}{\Gamma[1+\alpha+\beta]} \quad (17)$$

where we denoted $A \equiv K(1+\epsilon(1-x))$, $B \equiv K(1-\epsilon(1-x))$, $\alpha \equiv \frac{\tau_{N,+}}{\tau_{\xi,+}} = \frac{\lambda(1-\delta)}{1+\epsilon(1-x)}$, $\beta \equiv \frac{\tau_{N,-}}{\tau_{\xi,-}} = \frac{\lambda(1+\delta)}{1-\epsilon(1-x)}$, and $\Gamma[\alpha]$ is a Gamma function. Note that, for $\epsilon < 1$, normalization $Z_N(x)$ always exist, since in this case the behavior of the integrand near the endpoints behaves as $(A-N)^{\alpha-1}$ and $(N-B)^{\beta-1}$. These divergences are integrable for $\alpha, \beta > 0$, which are satisfied for $\epsilon < 1$.

For $1 - \epsilon(1-x) < 0$, the stationary pdf of tumor size, (16), is modified to

$$P_{\rm st}(N|x) = \frac{1}{Z_N(x)}\frac{1}{N^3}\left(\frac{K(1+\epsilon(1-x)) - N}{N}\right)^{\frac{\tau_{N,+}}{\tau_{\xi,+}}-1}\left(\frac{N + K(\epsilon(1-x)-1)}{N}\right)^{-\frac{\tau_{N,-}}{\tau_{\xi,-}}-1} \quad (18)$$

here $\tau_{N,-} \equiv \frac{1}{\epsilon(1-x)-1}$. Now the support of $P_{\rm st}(N|x)$ is $[0, K(1+\epsilon(1-x))]$. In contrast to previous case, here the integral can diverge on the support $[0, K(1+\epsilon(1-x))]$. Indeed, using the above introduced notations we have

$$Z_N(x) = \int_0^A \frac{1}{N^3}\left(\frac{A-N}{N}\right)^{\alpha-1}\left(\frac{N+B}{N}\right)^{-\beta-1} dN = \frac{A^{-1+\beta}B^{-1-\alpha}(A+B)^{-1+\alpha-\beta}(A\alpha + B\beta)\Gamma[\alpha]\Gamma[\beta-\alpha]}{\Gamma[1+\beta]} \quad (19)$$

where $B = K(\epsilon(1-x)-1)$ and $\beta = \frac{\lambda(1+\delta)}{\epsilon(1-x)-1}$, and the convergence of the integral requires $\alpha < \beta$. Indeed, the integrand diverges at zero as $\propto N^{-1-\alpha+\beta}$, and to have a integrable divergence at zero it is necessary to have $\beta - \alpha > 0$, note that both $\alpha, \beta > 0$. For biased environment toward bad state, that is $\delta < 0$, we get that the conditions $\alpha < \beta$, is satisfied for $x > 1 - \frac{1}{\epsilon|\delta|}$, that is just the opposite of the condition necessary for tumor extinction.

### IV. MORE ON PHASE SPACE PLAN OF THE EVOLUTIONARY OUTCOMES.

The exponents in the distributions (15) and (16) define the shape of the corresponding probability density. For the symmetric phenotype switching rates, that is $\mu_{EM} = \mu_{ME}$, and with no environmental bias, that is $\delta = 0$, we get that $\tau_{x,+} = \tau_{x,-} = 1/\sqrt{4\mu^2 + \epsilon^2}$ and $\tau_{\xi,+} = \tau_{\xi,-} = 1/\lambda$. While the characteristic timescale of the variation of tumor population size is $\tau_N \propto O((1-\epsilon)^{-1})$ which is of order of one for weak selection $\epsilon \propto 0$.

Let us consider the case, when $\tau_x > \tau_\xi$, that is environment changes more rapidly then the fraction of epithelial cells within the tumor. Here, we find that $\Pi_{st}(x \to x_g^*) \propto (x - x_g^*)^{\tau_x/\tau_\xi - 1} \to 0$, similarly for $\Pi_{st}(x \to x_b^*) \to 0$. Thus, once the environment fluctuates more rapidly than the fraction of epithelial cells, then it is highly unlikely observing the equilibrium state values of epithelial fraction corresponding to good or bad environments either.

When the environment fluctuates slower than the fraction of epithelial cells in the tumor, that is $\tau_\xi > \tau_x$, we observe the concentration of pdf function at the endpoints, that is $\Pi_{st}(x \to x_g^*) \propto (x - x_g^*)^{\tau_x/\tau_\xi - 1} \to \infty$ (still integrable).

Thus, we observe a noise-induced transition in the evolutionary outcome of the fraction of epithelial cells in the tumor.

Let us now examine the situation where $\tau_\xi = \tau_x$, that is the case where environmental fluctuations and the change of the composition of the tumor taking a place on the same timescale. This happens whenever $\lambda = \sqrt{4\mu^2 + \epsilon^2}$. Here again we get that in the absence of phenotype switching, that is $\mu = 0$, and in the environments where switching rate is equal to the selection pressure acting on the tumor cells, the distribution of the fraction of epithelial cells behaves as $\Pi_{st}(x) \propto 1/x(1-x)$, where the support is $x \in [0, 1]$. As it is seen from the distribution, the most probable values become either no epithelial cells or only epithelial cells in the tumor.

Therefore, for nonzero phenotype switching rate, $\mu \neq 0$, we get that the probabilities to observe tumors with the fraction of epithelial cells around either $x_g^*$ or $x_b^*$ are equal and given by, up to normalization constant, the following expression,

$$\Pi_{st}(x_b^*) = \Pi_{st}(x_g^*) \propto \frac{\epsilon^2}{4\mu^2\lambda^2}(\epsilon^2 - (2\mu - \lambda)^2) \tag{20}$$

In general, for the symmetric phenotype switching rates the possible pdfs of tumor population size and composition of it is presented in Fig.3.

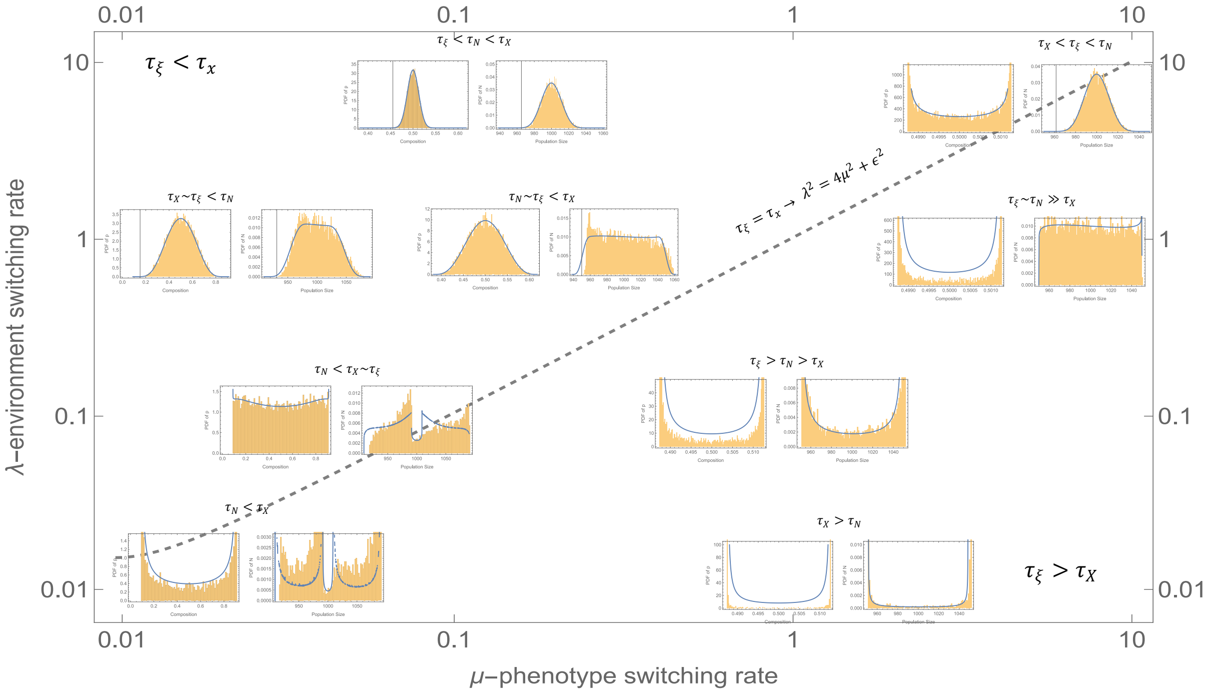


Supplementary Figure 3: Evolutionary outcomes of tumor size and composition, right and left subfigures in each pair, respectively, for various ordering of characteristic timescales. Bars show the pdfs obtained from the $10^4$ simulations with respective values of model parameters.

**IV.A. Fast-fluctuating environment: $\tau_\xi < \tau_X$, $M$-shape distribution.**

The extrema of the stationary probability density is found from (14) by taking the derivative of $\rho(z) = \rho(z, \xi = 1) + \rho(z, \xi = -1)$ over the state variable $z$. That is

$$\frac{d\rho}{dz} = \frac{2g(z)f(z)f'(z) - g'(z)\big(g(z)^2 + f(z)^2 + 2\lambda g(z)(f(z) + \delta g(z))\big)}{(g(z)^2 - f(z)^2)^2} e^{2\lambda \int^x \frac{f(z)+\delta g(z)}{g(z)^2 - f(z)^2}} \tag{21}$$

The equation of the extrema of the pdf is found by nullifying the derivative , that is $\frac{d\rho}{dz} = 0$ , which yields

$$f(z) + \delta g(z) + \frac{1}{\lambda} f'(z) f(z) - \frac{1}{2\lambda} g'(z) g(z) - \frac{1}{2\lambda} \frac{g'(z)}{g(z)} f(z)^2 = 0 \tag{22}$$

where $f(z)$ and $g(z)$ are introduced in (10).

For the fraction of epithelial cells in the tumor, these functions are $f(z) = \mu_{EM}(1 - z) - \mu_{ME} z$ and $g(z) = -\epsilon z(1 - z)$.

In general case, the left hand-side of (22) is a fifth-order polynomial expression for which the analytical solution is hard to find. Therefore, we focus here on the unbiased environment, $\delta = 0$, and assume symmetric switching rates $\mu_{EM} = \mu_{ME} = \mu$. For this case, we get the following solutions for (22)

$$z_1 = \frac{1}{2}, \tag{23}$$

$$z_{2,4} = \frac{1}{2}\big(1 \mp \sqrt{1 - \frac{4\mu}{\epsilon^2}(\lambda + \sqrt{\lambda^2 - \epsilon^2})}\big), \tag{24}$$

$$z_{3,5} = \frac{1}{2}(1 \mp \sqrt{1 + \frac{4\mu}{\epsilon^2}(\sqrt{\lambda^2 - \epsilon^2} - \lambda)}) \tag{25}$$

These are extrema for the probability density function of the fraction of epithelial cells. However, those extrema that are real must be in the support of $\Pi_{\rm st}(x)$, that is in the interval $[x_g^*, x_b^*]$. The equilibrium states $x_{b,g}^*$ are again symmetric with respect to 1/2. Namely, the equilibirum states are

$$x_{g,b}^* = \frac{1}{2}(1 \mp \zeta), \quad \zeta \equiv \frac{1}{\epsilon}(\sqrt{4\mu^2 + \epsilon^2} - 2\mu) \tag{26}$$

The extrema are all real, $z_i \in R \quad i = 1, ..5$, if $\lambda + \sqrt{\lambda^2 - \epsilon^2} \leq \frac{\epsilon^2}{4\mu}$ and $\lambda > \epsilon$. Therefore, $z_3$ and $z_5$ lie outside of the support in the fast-fluctuating regime $\tau_X > \tau_\xi$. Indeed, consider the following inequality $z_3 \leq x_g^* \leq z_2^*$. The last inequality holds whenever

$$\sqrt{1 + \frac{4\mu}{\epsilon^2}(\sqrt{\lambda^2 - \epsilon^2} - \lambda)} \geq \zeta \geq \sqrt{1 - \frac{4\mu}{\epsilon^2}(\sqrt{\lambda^2 - \epsilon^2} + \lambda)} \tag{27}$$

Simplifying the last expression, we get

$$\lambda - \sqrt{\lambda^2 - \epsilon^2} < \sqrt{4\mu^2 + \epsilon^2} - 2\mu < \lambda + \sqrt{\lambda^2 - \epsilon^2} \tag{28}$$

The fast-fluctuating regime, $\tau_X > \tau_\xi$, implies that $\lambda > \sqrt{4\mu^2 + \epsilon^2}$. In this case, the both inequality in (28) always satisfied, and points $z_3$, $x_g^*, z_2$ merge if $\lambda = \sqrt{4\mu^2 + \epsilon^2}$. Note, that in the fast-fluctuating regime, the stationary pdf vanishes in the boundaries, that is $\Pi_{\rm st}(x) \to 0$ for $x \to x_{g,b}^*$. Thus, the only possibility in this case, is $z_1 = \frac{1}{2}$ is minima, and $z_2, z_4$ are symmetrically located maxima around $z_1$. In case of slow-environment, that is $\lambda < \sqrt{4\mu^2 + \epsilon^2}$, we can get inverted $M$-shape distribution. This happens, whenever $\lambda > \epsilon$ and $\lambda - \sqrt{\lambda^2 - \epsilon^2} < \frac{\epsilon^2}{4\mu}$. In this case, though, $z_2$ and $z_4$ are complex, and $z_3$ and $z_5$ are shifted into the interval $[x_g^*, x_b^*]$.

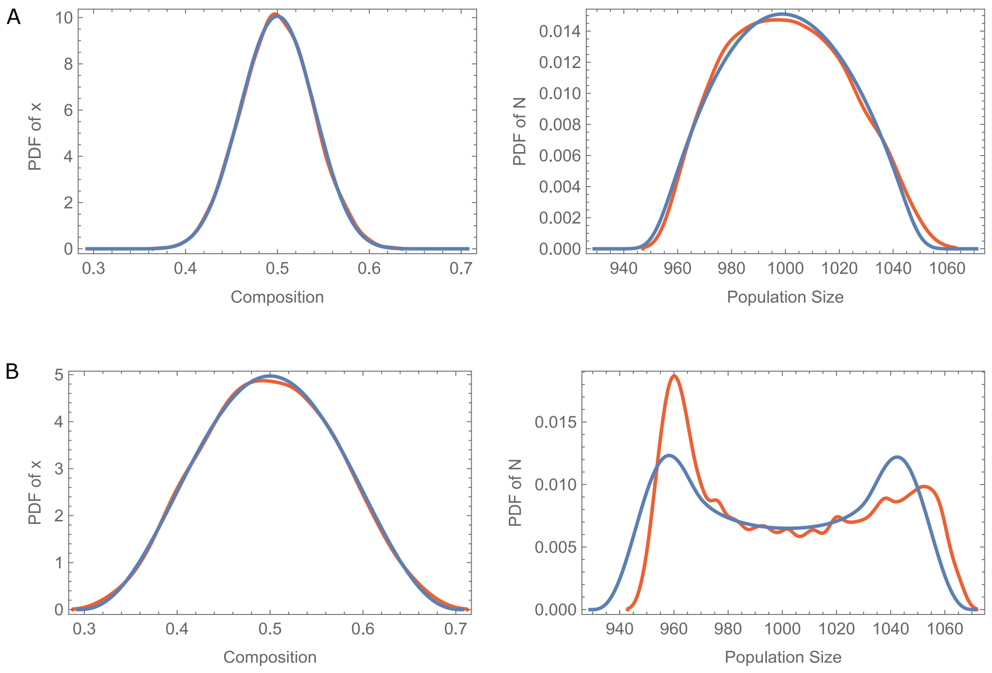


Supplementary Figure 4: Evolutionary outcomes of tumor size and composition, right and left subfigures in each pair, respectively, for fast fluctuating environment.
A). Pdf of tumor composition and size, left and right subfigures, respectively, for $\tau_X > \tau_N > \tau_\xi$. Here model parameters are $\mu = 0.05, \lambda = 2$.
B). Pdf of tumor composition and size for $\tau_X > \tau_\xi > \tau_N$, with model parameters $\mu = 0.05, \lambda = 0.5$. The remaining model parameters are same for both cases and are as follows $\epsilon = 0.1, \delta = 0, K = 1000$. The red lines show the pdfs obtained from $10^4$ simulation runs with respective model parameters.

In the limiting case of fast-fluctuating environment, that is $\lambda \to \infty$, we get that the most probable value of the pdf of fraction of epithelial cells is given by

$$x_{\text{mpv}} = \frac{1}{2\epsilon\delta}\left((\mu_{EM} + \mu_{ME} + \epsilon\delta) \mp \sqrt{(\mu_{EM} + \mu_{ME} + \epsilon\delta)^2 - 4\epsilon\delta\mu_{EM}}\right) \tag{29}$$

, as one may get by replacing the stochastic term in (10) by its average value. Note, that $x_{\text{mpv}} \in [0, 1]$ which is satisfied only by one of the possible values given from (29).

Depending on the characteristic timescale differences between the evolution of tumor size and composition, it is possible to obtain two qualitatively different regimes, beyond $M$-shape distribution, that are presented in Fig.4.

The pdfs of tumor size and composition are unimodal when the environment fluctuates faster than both of them, that is for $\tau_\xi < \tau_N < \tau_X$ and $\tau_\xi < \tau_X < \tau_N$. The unimmodality of the tumor size is due to (16), where for each possible value of $x$ the pdf is concentrated between the endpoints, since the exponents are negative.

Therefore, pdf support for tumor size and composition behaves differently, as it is illustrated in the main text, in $\tau_\xi < \tau_N < \tau_X$ and $\tau_\xi < \tau_X < \tau_N$ cases. Indeed, the length of the support of $\Pi_{st}(x)$ is equal to $x_b^* - x_g^* = \frac{1}{\epsilon}(\sqrt{4\mu^2 + \epsilon^2} - 2\mu)$. For fixed selection strength $\epsilon = \text{const}$ the condition $\tau_\xi < \tau_X < \tau_N$ is achieved by decreasing $\tau_X = \frac{1}{\sqrt{4\mu^2+\epsilon^2}}$ which amounts to an increased phenotype switching rate $\mu$, resulting in shrinkage of the support $x_b^* - x_g^*$.

A qualitatively different phenomenon occurs in cases when the tumor size evolves much faster than the environment and the fraction of the epithelial cells within the tumor $\tau_N < \tau_\xi < \tau_X$. In this case, tumor size evolves as in the fixed environment either in $\xi = 1$ or $\xi = -1$. Now, in the fixed environment, the size of the tumor evolves along the variations in the tumor composition $x(t)$, that takes place in much longer times than that of the environment, hence it evolves towards the vicinity of the most probable value $x_{\text{mpv}}$ (see left subfigure in Fig.4B). Therefore, the tumor sizes has two peaks that are given by $K(1 \pm \epsilon(1 - x_{\text{mpv}}))$, as it can be obtained from (16) by noting that $\tau_N < \tau_\xi$.

#### IV.B. Slow environment: $\tau_X < \tau_\xi$.

In contrast to the fast-fluctuating environment, in the slow environment, the pdf of tumor composition is bimodal with peaks on the endpoints $\left\{x_g^*, x_b^*\right\}$, as follows from (15) for $\tau_X < \tau_\xi$.

Nevertheless, it is still possible to observe a uniform and almost flat distribution of tumor size due to the difference in characteristic timescales of the underlying processes, though the time-evolution of the tumor size is affected by the evolution of tumor composition and not vice versa.

The unimodal size distribution of the tumor can be observed in cases where $\tau_X < \tau_\xi < \tau_N$ (see Fig.3A upper right). Here, the fraction of epithelial cells relaxes to its equilibrium value in the current environment before the switch occurs, on average. Due to the last observation, in (5), we can substitute $\xi(t)(1-x(t)) \to \frac{1-x_g^*}{2} - \frac{1-x_b^*}{2} = \frac{1}{2\epsilon}(\sqrt{4\mu^2 + \epsilon^2} - 2\mu)$. The tumor size will then evolve to $\hat{N} = K(1 + \frac{1}{2}(\sqrt{4\mu^2 + \epsilon^2} - 2\mu))$

Therefore, for given $\epsilon$, the composition of the tumor will evolve faster than the size of the tumor population, that is, $\tau_X < \tau_N$ will be fulfilled if the dynamics of the tumor composition is driven primarily by the phenotype switching process rather than selection resulting into shrinkage of the

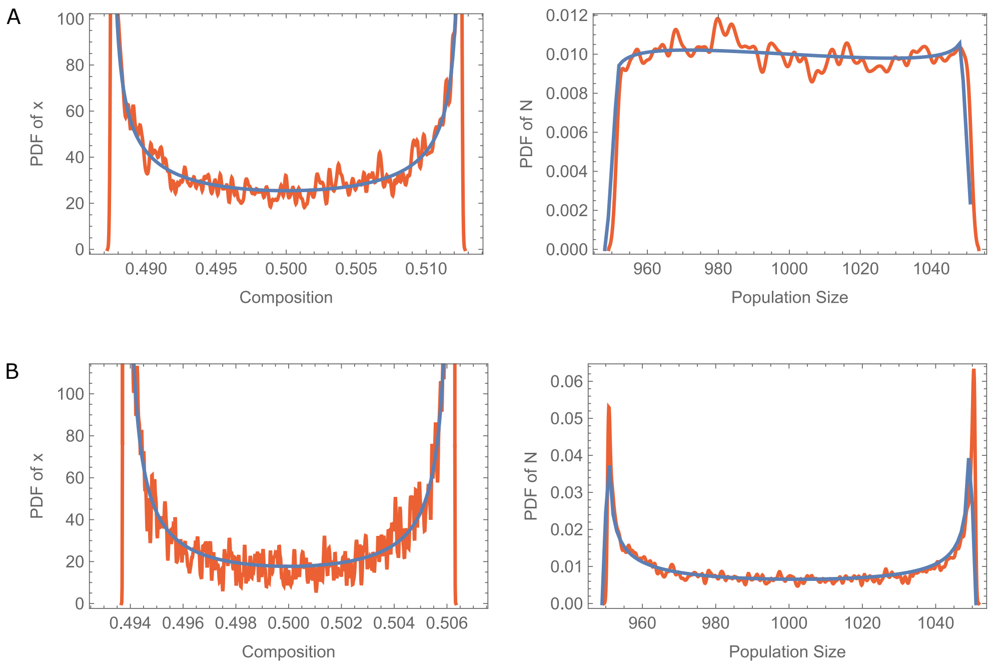


Supplementary Figure 5: Evolutionary outcomes of tumor size and composition, right and left subfigures in each pair, respectively, for slow-fluctuating environment.
A). Pdf of tumor composition and size, left and right subfigures, respectively, for $\tau_\xi \sim \tau_N > \tau_X$. Here model parameters are $\mu = \lambda = 1$.
B). Pdf of tumor composition and size for $\tau_\xi > \tau_N > \tau_X$, with model parameters $\mu = 2, \lambda = 0.5$. The remaining model parameters are same for both cases and are as follows
$\epsilon = 0.1, \delta = 0, K = 1000$. The red lines show the pdfs of relevant quantities through smoothing the histogram obtained from $10^4$ simulation runs with respective model parameters.

support of $\Pi_{st}(x)$, Fig.3A upper right corner.

In case of comparable $\tau_N$ and $\tau_\xi$ (see Fig.5B), the above procedure is not justified, since substituting fluctuating part of the tumor size dynamics by its average is not allowed. Therefore, we can get an estimate of $P_{st}(N)$ in this case as well.

Indeed, when $\tau_N \approx \tau_\xi$, then from (16) we get that $Z_N(x) \approx \frac{2\epsilon(1-x)}{K^2}$. Inserting into $P_{st}(N) = \int_{x_g^*}^{x_b^*} \Pi(x) P_{st}(N|x)dx$, we obtain that

$$P_{st}(N) \approx \frac{K^2}{2\epsilon}\frac{1}{N^3}(1 - \int_{x_g^*}^{x_b^*} x\Pi_{st}(x)dx) \approx \frac{K^2}{2\epsilon}\frac{1}{N^3}(1 - \frac{x_b^* - x_g^*}{2}) \approx \frac{K^2}{2\epsilon}\frac{1}{N^3} \tag{30}$$

where we used the fact that $\Pi_{st}(x)$ is symmetric and concentrated in the endpoints, thus the average is substituted by the difference. For the considered values of model parameter we get that $P_{st}(N) \sim 10^{-2}$. Note, that $P_{st}(N)$ is defined in the range $\big\{K(1-\epsilon(1-x_g^*)), K(1+\epsilon(1-x_g^*))\big\}$.

When environment evolves slowly than the dynamics of the size of tumor, $\tau_\xi > \tau_N > \tau_X$, the size distribution becomes bimodal as well, since the exponents become negative in (16). Indeed, in this case both tumor size and composition evolve to their equilibrium values in the given environment before environment switches. Then the average population size can be defined by the average over the environmental fluctuations of the equilibrium values that are given by $N_b^* = K(1-\epsilon(1-x_b^*))$ and $N_g^* = K(1+\epsilon(1-x_g^*))$ in bad and good environment, respectively. Therefore, in this case the shape of the tumor size distribution is bimodal, (see Fig.5B).

In the biased environment, the above considerations hold for the cases $\tau_\xi > \tau_{N\pm}$ and $\tau_{N\pm} > \tau_\xi$ as well, since the tumor composition evolves much faster in both good and bad environments, and the average tumor size is given by

In case of slow variation of the tumor composition w.r.t the size evolution, that is $\tau_\xi > \tau_X > \tau_N$, we can make progress by considering only the diagonal terms in $P_{st}(N) = \int_{x_g^*}^{x_b^*} \Pi(x) P_{st}(N|x)dx$. That is, decomposing the last expression into the following terms

$$P_{st}(N) = \sum_{\xi=\pm1} P_{st}(N,\xi|x)\Pi_{st}(x) = \sum_{\xi=\pm1}\sum_{\xi'=\pm1} P_{st}(N,\xi|x)\Pi_{st}(x,\xi'=\pm1) \tag{31}$$

$$= \sum_{\xi=\pm1} P_{st}(N,\xi|x)\Pi_{st}(x,\xi) + \sum_{\xi=\pm1} P_{st}(N,\xi|x)\Pi_{st}(x,-\xi) \tag{32}$$

Now, if we keep only the diagonal term (first term in (31)), for large characteristic times of we will get more and more concentrated, that is, the probability of observing tumor of size $K(1-\epsilon(1-x_b^*))$ and $K(1+\epsilon(1-x_g^*))$ decreases, see Fig.6.

Here, we should mention that for $\tau_\xi > \tau_N > \tau_X$ we must also get the discrepancy, because both distributions (15) and (16) are bimodal. However, we do not observe the discrepancy, since for $\tau_\xi > \tau_N > \tau_X$ the distance between the neighboring peaks of $P_{st}(N|x)$ is $|K(1-\epsilon(1-x_g^*)) - K(1+\epsilon(1-x_b^*))| \sim \epsilon K(x_b^* - x_g^*) = K(\sqrt{4\mu^2+\epsilon^2} - 2\mu) \approx \epsilon^2 K/4\mu$. In the last derivation we expand $\sqrt{4\mu^2+\epsilon^2}$ w.r.t $\epsilon$, since $\tau_X = \frac{1}{\sqrt{4\mu^2+\epsilon^2}} < \tau_N = O(1\pm\epsilon)$. So far we have considered the case of unbiased environment $\delta = 0$ and symmetric phenotype switching rates $\mu_{EM} = \mu_{ME} = \mu$. In the general case, the distributions are again either unimodal (it can be peaked on the edge, in contrast to symetric case where it is not possible) or bimodal, however, they are not symmetric anymore. The analysis for the biased environment and asymmetric phenotype switching rates is carried out analogously, that is, figuring out the ordering of the timescale differences, then considering the

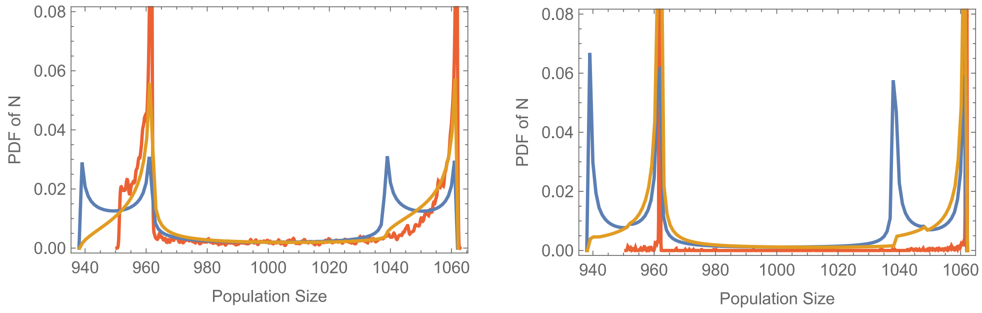


Supplementary Figure 6: Tumor size distribution in the regime of slow-fluctuating environment, $\tau_\xi > \tau_X > \tau_N$, for varios $\tau_\xi$.
The left and rigth panels correspond to $\lambda = 0.1$ and $\lambda = 10^{-3}$, respectively. The blue and orange lines on the right panel show the pdfs of tumor size obtained from $P_{\rm st}(N)$ by keeping and disregarding non-diagonal terms in the Bayes rule, see (31). The red lines show the pdf of respective quantities obtained from the outcomes of simulations. Here the remaining model parameters are $\epsilon = 0.1, \mu = 0.1, \delta = 0, K = 1000$.

stationary distributions (15 and (16).

### V. LONG-TERM GROWTH RATE

Using (15) and (16) one may obtain the average population size of the tumor after transient period is passed. The average population size is given by

$$\langle N \rangle = \int_{x_g^*}^{x_b^*} dx \Pi_{\rm st}(x) \int_{K(1-\epsilon(1-x))}^{K(1+\epsilon(1-x))} N P_{\rm st}(N|x) dN \tag{33}$$

Using the notations in (17), the integral over the population size $N$ is evaluated to

$$\int_{K(1-\epsilon(1-x))}^{K(1+\epsilon(1-x))} N P_{\rm st}(N|x) dN = \frac{1}{Z_N(x)} \int_B^A \frac{1}{N^2} \left(\frac{A-N}{N}\right)^{\alpha-1} \left(\frac{N-B}{N}\right)^{\beta-1} = \tag{34}$$

$$= \frac{1}{Z_N(x)} \frac{A^{-\beta} B^{-\alpha} (A-B)^{\alpha+\beta-1} \Gamma[\alpha]\Gamma[\beta]}{\Gamma[\alpha+\beta]} = \frac{AB(\alpha+\beta)}{\alpha A + \beta B} = K(1 + \epsilon\delta(1-x)) \tag{35}$$

Inserting (34) back into (33) we get

$$\langle N \rangle = K(1 + \epsilon\delta(1 - \langle x \rangle)) \tag{36}$$

, where $\langle x \rangle = \int_{x_g^*}^{x_b^*} x \Pi_{\rm st}(x) dx$. From the last relation it follows that in an unbiased environment $\delta = 0$, the average population size is always equal to $K$. Therefore, we already observe a case, where

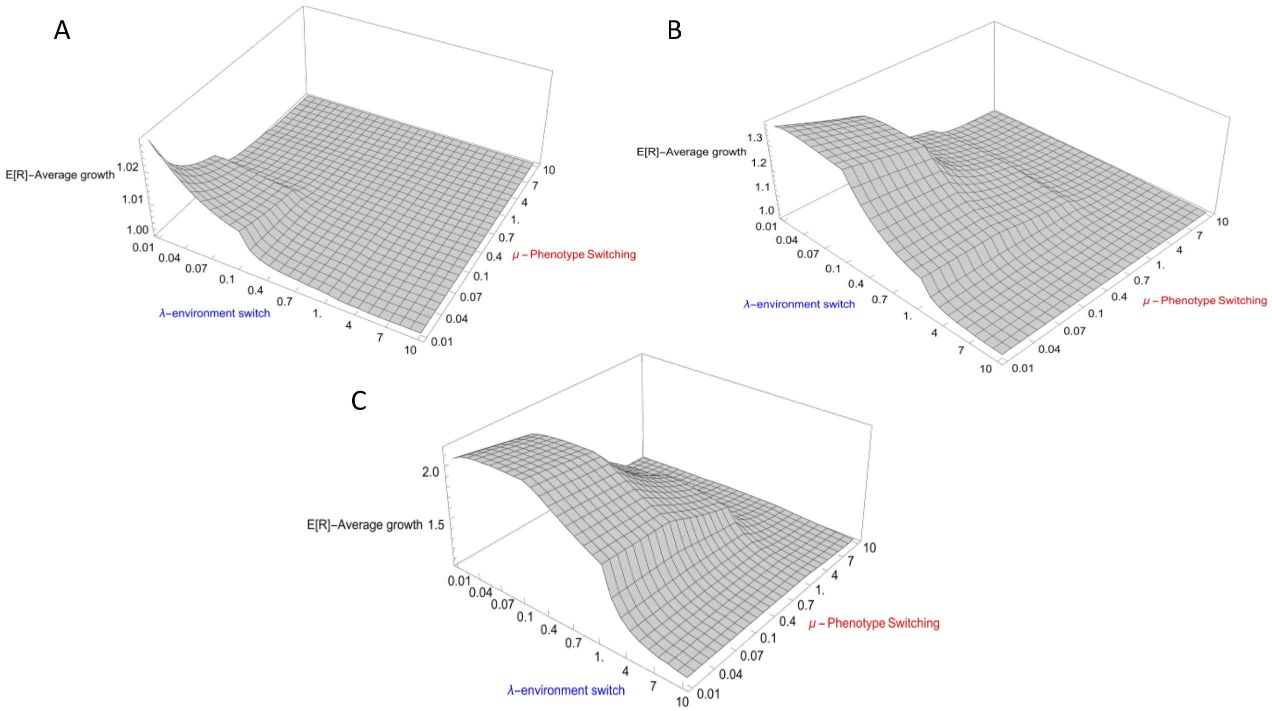


Supplementary Figure 7: Long-term growth rate of the tumor, (37), evaluated for different values of environment switching rates, selection advantage and phenotype switching rates. The phenotype switching is assumed to be symmetric, $\mu_{EM} = \mu_{ME} = \mu$. Both $\mu$ and $\lambda$ are evaluated at points $[0.01, 0.1] \cup [0.2, 1] \cup [2, 10]$ with step size 0.01, 0.1 and 1 for each interval, respectively. A), B) and C) are long-term growth rate for $\epsilon = 0.01$, $\epsilon = 1$, and $\epsilon = 3$, respectively.

even in the unbiased environment and symmetric phenotype switching rates the size distribution is not symmetric (see Fig.4B).

A better approximation is first defining the average growth rate of the population, then approximate the average size of the tumor by this averaged growth rate. The growth rate of the population is $R(\xi, x) = 1 + \epsilon\xi(t)(1 - x(t))$, now assuming that $x(t)$ is in its stationary state we can approximate the growth rate by

$$
\begin{aligned}
E_{X,\xi}[R] &= \sum_{\xi=\pm 1} \int_{x_g^*}^{x_b^*} R(\xi, x)\Pi_{st}(x,\xi)dx \qquad (37)\\
&= \int_{x_g^*}^{x_b^*} R(\xi=1,x)\Pi_{st}(x,\xi=1)dx + \int_{x_g^*}^{x_b^*} R(\xi=-1,x)\Pi_{st}(x,\xi=-1)dx = \\
&= 1+\epsilon\int_{x_g^*}^{x_b^*} (\Pi_{st}(x,\xi=1)-\Pi_{st}(x,\xi=-1))dx - \epsilon\int_{x_g^*}^{x_b^*} x(\Pi_{st}(x,\xi=1)-\Pi_{st}(x,\xi=-1))dx = \\
&= 1+\epsilon\delta+\epsilon\int_{x_g^*}^{x_b^*} x(\Pi_{st}(x,\xi=-1)-\Pi_{st}(x,\xi=1))dx \qquad (38)
\end{aligned}
$$

Here we used the relation $\delta = \int_{x_g^*}^{x_b^*}(\Pi_{st}(x,\xi=1)-\Pi_{st}(x,\xi=-1))dx$. Using (37), the average population size is then approximated as $N^* = E_{x,\xi}[R]K$. Note that even in an unbiased environment, $\delta = 0$, the average size of the tumor is not equal to the carrying capacity of the environment, $N^* \neq K$, in general, because $E[R] \neq 1$ due to the last term in (37). The dependency of the long-term growth rate on the model parameters is illustrated in Fig. 8. It increases with increasing selective advantage, $\epsilon$. While decreases with increasing phenotype switching rates and the rate of environmental fluctuations. Here we consider symmetric phenotype switching rates. The long-term growth rate is bigger for small environmental and phenotypic switching rates, regardless of the selection strength $\epsilon$. Therefore, for the small selection strength no phenotype switching is beneficial for the population, see Fig.7A. Increasing $\epsilon$ results into the shift of maximum of the long-term growth rate towards the interior of the considered intervals of environment switching rates, $\lambda$, and phenotype switching rate, $\mu$.

The average population size obtained from (37) agrees with the simulation results, in all orderings of characteristic timescales, except in $\tau_\xi > \tau_X > \tau_N$.

Indeed, in this case, the simulation results and the prediction obtained through (37) reveal a different trend of the average population size w.r.t the selection strength $\epsilon$, see Fig.8A. The prediction from (37) is that increasing the selection advantage/disadvantage of $M$-cells in a biased environment towards the bad state, $\delta = -0.5$, will decrease the average tumor size. However, in slow-environment, simulation reveal that opposite holds (see Fig.8A).

Indeed, this effect is explained by noticing that first the approach that leads to (37) fails in slow environment, because averaging over environment is not "allowed". The average population size in the quenched noise limit, that is the stochasticity of the environment is through the initial state only, is given by

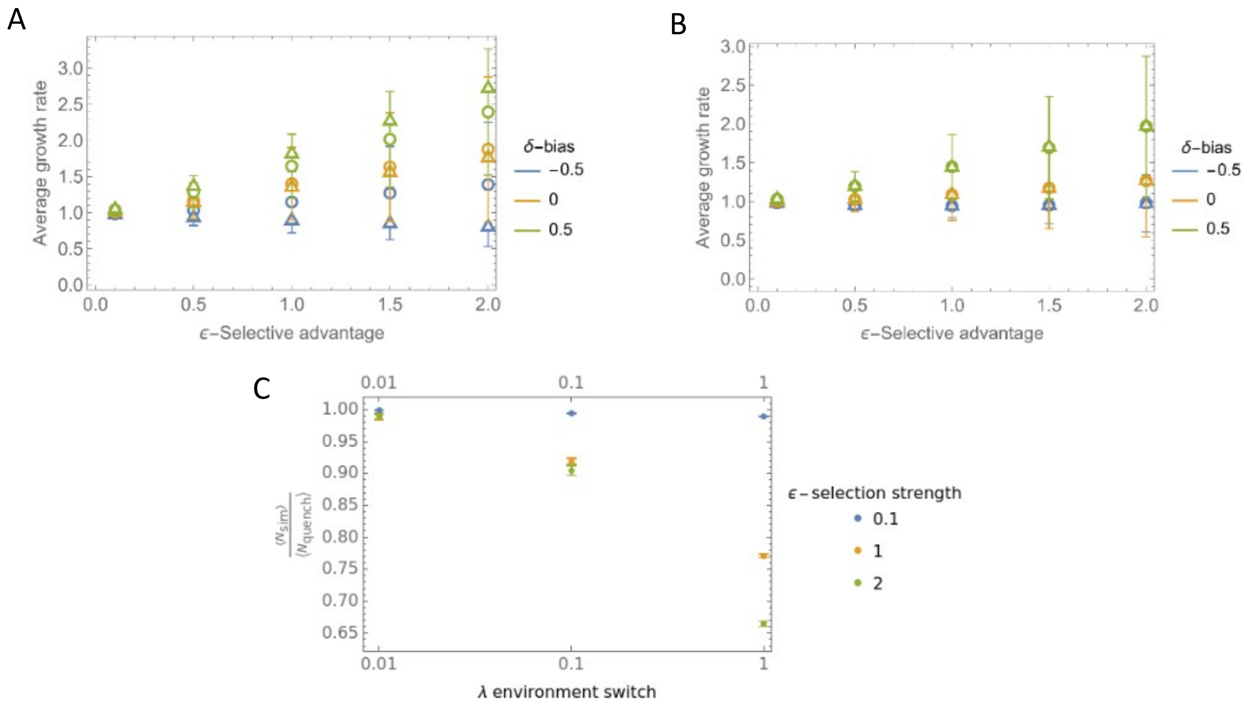


Supplementary Figure 8: Average growth rate of the population obtained from (37) and simulations for various model parameters. A) and B) average growth rate of the population for slow and fast environments $\lambda = 10^{-2}$ and $\lambda = 1$, respectively. Circles and triangles show growth rate obtained from simulations and from evaluation of long-term growth rate, (37). In each figure $\delta$ is the bias of the environment.

C) Comparison of the average population size in the quenched disorder limit, (39), and simulation results for various $\epsilon$ and $\lambda$, for unbiased environment, $\delta = 0$. The remaining model parameters are $\mu_{EM} = \mu_{ME} = 0.1$ and $K = 10^3$.

$$N_{\text{quench}} = K(1 - \epsilon\delta + \epsilon(\frac{1+\delta}{2}x_b^* - \frac{1-\delta}{2}x_g^*)) \tag{39}$$

Using (39), we find that $N^*/K > 1$, that is the tumor average population size is greater than carrying capacity $K$, if $|\delta| < 2x_b^* - 1$ for given $\delta < 0$, we assumed symmetric phenotype switching rates $\mu_{EM} = \mu_{ME} = \mu$. Thus, the biased environment towards bad state still allows the tumor to growth, as it is revealed in the simulation (see Fig.8A).

On the other hand, the minimum of $N^*$ is reached at point $\epsilon_m = \frac{2\mu|\delta|}{1-\delta^2}$.That is for given bias $\delta$, there is an $\epsilon$ selection strength that impedes tumor growth.

Therefore, we observe that phenotype switching does not provide an advantage in this case, since the average population size will be greatest whenever $1 + \epsilon\delta + \frac{\epsilon x_b^*}{2}(1-\delta) - \frac{\epsilon x_g^*}{2}(1+\delta)$ is

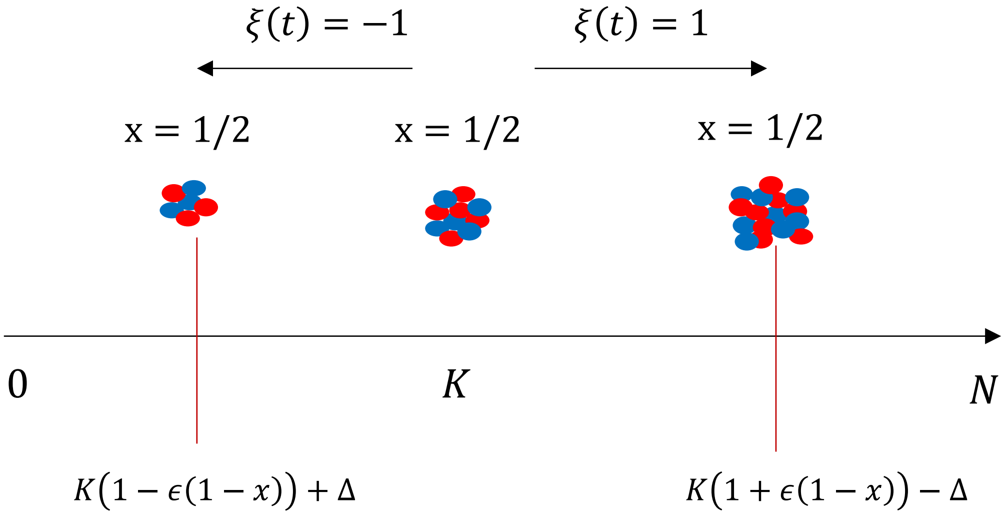


Supplementary Figure 9: Graphical representation of the appearance of the worse outcome for a fixed fraction of epithelial cells in the tumor. In a good environment, population size increases and hits the right wall located at $K(1+\epsilon(1-x)-\Delta$, where $\Delta \ll K$. In a bad environment, population size decreases and eventually hits the wall located at $K(1-\epsilon(1-x)+\Delta$

.

maximized over $\mu_{EM}$ and $\mu_{ME}$, which is maximized if $x_b^* = 1$ and $x_g^* = 0$, that happens whenever $\mu_{EM} = \mu_{ME} = 0$.

Note, that the discrepancy between simulation and numerical evaluation of (37) vanishes with increasing environmental switching rate, $\lambda$, see Fig.8B. In this case (39) does not correctly predicts the average population size, see Fig.8C. The discrepancy increases with increasing environmental switching rate $\lambda$, that is the quenched disorder limit is not valid for increased $\lambda$.

The phenotype switching is beneficial in the opposite case, that is, $\lambda \to \infty$, Therefore, there is no balance between the phenotype switching rates in the biased but fast-fluctuating environment. In this regime using (29), we can approximate the long-term growth rate of the tumor by $1+\epsilon\delta(1-x_{\mathrm{mpv}})$. Maximizing over phenotype switching rates we get that in bad and good environment, $\delta < 0$ and $\delta > 0$, the frequency of $E \to M$ switching is $\frac{\mu_{em}^*}{\mu_{em}^*+\mu_{me}^*} \to 1$ or $\frac{\mu_{em}^*}{\mu_{em}^*+\mu_{me}^*} \to 0$.

### VI. HITTING PROBABILITITIES AND HITTING TIMES

The first hitting probability under dichotomous noise is widely discussed for a variety of models [2–4]. As it is mentioned in the main text, the worse case outcome of the tumor population size is $\mathrm{Max}[K(1-\epsilon(1-x)), 0]$ for the fixed composition of the tumor, $x$. We are interested in the probability of the occurrence of the worse outcome. The graphical representation of the process is presented in Fig.9. For a fixed fraction of epithelial cells the good environment

To obtain the probability of eventually hitting the left wall first, instead of right one, we assume that these walls are absorbing. Hence, for the fixed composition of epithelial cells, $x$, the tumor population size eventually attains one of the possible values at the boundaries, that is $K(1 \pm \epsilon(1-x))$.

Let us denote by $\psi_+(N_0, x, t)$ and $\psi_-(N_0, x, t)$ the probability that the tumor population size attains the minimal value before or at time $t$, that is $N(t) = K(1-\epsilon(1-x)) + \Delta$, with $N_0$ initial population size and $\xi = \pm 1$ initial state of the environment. We assume that the fraction of epithelial cells is fixed $x = \text{const}$.

If the process begins at $\xi = 1$, then in the small time interval $dt$ either environment remains fixed, hence population size evolves in the $\xi = 1$ state, or environment switches to $\xi = -1$ state, and the population evolves in $\xi = -1$ state. Hence, the hitting probability $\psi_+(N_0, x, t)$ satisfies the following relation

$$\psi_+(x, N_0, t+dt) = (1-\lambda dt)\psi_+(x, N_0 + (f(N_0) + g(N_0))dt, t) + \lambda dt\ \psi_-(x, N_0 + (f(N_0) - g(N_0))dt, t), \tag{40}$$

$$\psi_-(x, N_0, , t+dt) = (1-\lambda dt)\psi_-(x, N_0 + (f(N_0) - g(N_0))dt, t) + \lambda dt\ \psi_+(x, N_0 + (f(N_0) + g(N_0))dt, t) \tag{41}$$

where $f(N) = N(1-\frac{N}{K})$ and $g(N) = \epsilon(1-x)N$. Expanding (40) and (41) for small $dt$ and neglecting the terms $\propto dt^2$, we get

$$\partial_t \psi_+(x, N, t) = \big(f(N) + g(N)\big)\partial_N \psi_+(x, N, t) - \lambda \psi_+(x, N, t) + \lambda\ \psi_-(x, N, t), \tag{42}$$

$$\partial_t \psi_-(x, N, t) = \big(f(N) - g(N)\big)\partial_N \psi_-(x, N_0, t) + \lambda \psi_+(x, N, t) - \lambda\ \psi_-(x, N, t), \tag{43}$$

where we substitute $N_0 \to N$, since the initial state is arbitrary. We are interested in $\lim_{t\to\infty} \psi_\pm(x, N, t) = \psi_\pm(x, N)$, hence we get the left hand-side nullifies. Denoting $\psi(N, x) = \psi_+(N, x) + \psi_-(N, x)$ and $\varphi(N, x) = \psi_+(N, x) - \psi_-(N, x)$ we get from (42) and (43) the following pairs of differential equations

$$f(N)\partial_N \psi \ = \ -g(N)\partial_N \varphi, \tag{44}$$

$$g(N)\partial_N \psi + f(N)\partial_N \varphi \ = \ 2\lambda \varphi \tag{45}$$

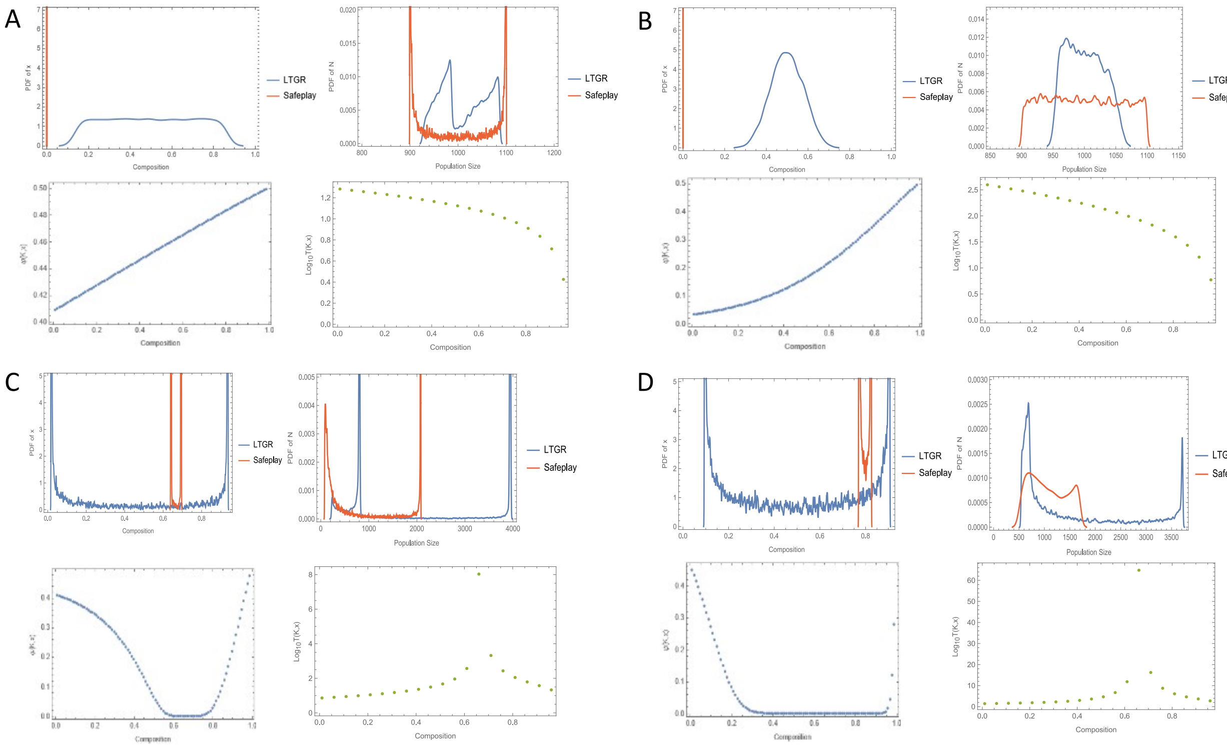


Supplementary Figure 10: Evolutionary outcomes for the optimal values of phenotype switching rates that minimize the probability of the worst case outcome or maximize the survival time. In each panel, the blue and red curves show pdf obtained from the simulation with optimal phenotype switching rates that maximize the long-term growth rate and survival time, respectively. The right and left sub-panel in the second row in each panel show the hitting probabilities and survival times for various fraction of epithelial cells in the tumor,$x$. In A),B) C) and D) the model parameters are $\epsilon = 0.1,\ \lambda = 0.1$, $\epsilon = 0.1,\ \lambda = 1$, $\epsilon = 3,\ \lambda = 0.1$ and $\epsilon = 3,\ \lambda = 1$, respectively. The remaining model parameters are $\delta = 0$ and $K = 1000$ and $\Delta = 1$. The initial size of the tumor is randomly selected in the simulations from $[\mathrm{Max}[K(1-\epsilon(1-x)), 0], K(1+\epsilon(1-x))]$. The pdfs are obtained from $10^4$ independent simulations. The values of $x$ and $N$ are taken after $100\mathrm{Max}[\tau_{\mathrm{x},\pm}, \tau_{\xi,\pm}, \tau_{\mathrm{N},\pm}]$

From (44) and (45), we find that $\varphi(N,x) = e^{\int_{K(1+\epsilon(1-x))-\Delta}^{N} \frac{2\lambda f(z)}{f^2(z)-g^2(z)}dz}$, and for the hitting probability we find

$$\psi(x,N) = \frac{1}{\mathcal{Z}} \int_{K(1+\epsilon(1-x)-\Delta}^{N} \frac{2\lambda g(z)}{g^2(z)-f^2(z)} \varphi(x,z)dz \tag{46}$$

where $\mathcal{Z}$ is the normalization constant. It is found by considering the boundary conditions,

$\psi_+(x, K(1+\epsilon(1-x)) - \Delta) = 0$ and $\psi_-(x, K(1-\epsilon(1-x)) + \Delta) = 1$. These conditions are satisfied for $\mathcal{Z} = \int_{K(1+\epsilon(1-x)-\Delta}^{,K(1-\epsilon(1-x))+\Delta} \frac{2\lambda g(z)}{g^2(z)-f^2(z)}\varphi(x,z)dz - \Delta) - \varphi(K(1-\epsilon(1-x)) + \Delta) - 1$.

Simplifying the integrant in (46) we find that

$$\psi(n,x) = \frac{1}{\mathcal{Z}}(\lambda \int_{K(1+\epsilon(1-x))-\Delta}^{n} (\frac{1}{\epsilon z(1-x) - z(1-z/K)} + \frac{1}{\epsilon z(1-x) + z(1-z/K)})\phi(z)dz - 1) \quad (47)$$

In the main text the initial state of the population is assumed to be $N = K$.

To find the survival time of the tumor before the worse outcome happens, which can be extinction for $\epsilon > 1$, we substitute the absorbing boundary at $K(1+\epsilon(1-x)) - \Delta$ by reflective boundary [3, 5, 6].

The differential equation governing the survival time of tumor are as follows

$$(f(N) + g(N))\partial_N \tau_+ - \lambda\tau_+ + \lambda\tau_- = -1, \quad (48)$$

$$(f(N) - g(N))\partial_N \tau_+ + \lambda\tau_+ - \lambda\tau_- = -1 \quad (49)$$

where $\tau_\pm$ is the mean first hitting with initial state $\xi = \pm 1$.

We are interested in the unconditional hitting times, that is $\tau = \frac{1}{2}(\tau_+ + \tau_-)$. To find the hitting time $\tau$, we represent (48) as follows

$$\mathcal{L}_+\tau_+ = -1 - \lambda\tau_-, \quad (50)$$
$$\mathcal{L}_-\tau_- = -1 - \lambda\tau_+,$$

where $\mathcal{L}_\pm = (f \pm g)\partial_N - \lambda$ is an operator acting on the quantities right from it.

Acting by $\mathcal{L}_-$ and $\mathcal{L}_+$ on the first and second equations of (50), and using (50) we get

$$(f^2 - g^2)\partial_N^2\tau_+ + ((f-g)(\partial_N f + \partial_N g) - 2\lambda f)\partial_N\tau_+ = 2\lambda, \quad (51)$$
$$(f^2 - g^2)\partial_N^2\tau_+ + ((f+g)(\partial_N f - \partial_N g) - 2\lambda f)\partial_N\tau_+ = 2\lambda$$

Summing the equations in (51), and noting that $\partial_N(\tau_+ - \tau_-) = \frac{-2-f\partial_N\tau}{g}$, we get for $\tau(N)$ the following differential equation

$$(g(N)^2 - f(N)^2)\partial_N^2\tau + \big((g(N)\partial_N g(N) - f(N)\partial_N f(N) + 2\lambda f(N) + \frac{f(N)}{g(N)}(f(N)\partial_N g(N) \quad (52)$$
$$-g(N)\partial_N f(N)\big)\partial_N\tau = -2\lambda + \frac{g(N)\partial_N f(N) - f(N)\partial_N g(N)}{g(N)}$$

Solving (52) numerically with boundary conditions $\partial_N \tau_N|_{K(1+\epsilon(1-x))-\Delta} = 0$ and $\tau(\max[0, K(1-\epsilon(1-x))] + \Delta) = 0$ yields the behavior of hitting times. In Fig.10 the behavior of hitting times and hitting probabilities are presented for different values of fraction of epithelial cells in the tumor,$x$.

As it is seen from the figure, both hitting probabilities and survival times attain their maximum values at $\hat{x} = \mathrm{Max}[0, 1 - \frac{1}{\epsilon}]$. We analyze the behavior of the mean hitting times for sticky boundary conditions as well, that is Eq.(48) with the following boundary conditions $\tau_+|_{K(1+\epsilon(1-x))-\Delta} = \tau_-|_{K(1+\epsilon(1-x))-\Delta} + 1/\lambda$. In this case, the tumor close sticks on the right boundary till the environment switches to bad state, the waiting time till the environment switches is $1/\lambda$ on average. Therefore, the optimal fraction of epithelial cells is again $\hat{x} = \mathrm{Max}[0, 1 - \frac{1}{\epsilon}]$.

Indeed, the optimal fraction of epithelial cells, for any $\epsilon$, is given by $\hat{x} = \mathrm{Max}[0, 1 - \frac{1}{\epsilon}]$. The worst case outcome for the tumor can happen in the bad environment, $\xi = -1$ in (3). The time needed to hit the absorbing state $N_b^* + \Delta$ is inversely proportional to the right hand side of (3), that is the velocity of the process. Considering the behavior of the time needed to hit the absorbing state started from the vicinity of the absorbing state one gets it scales as $\sim |1 - \epsilon(1-x)|^{-1}$, which diverges at the optimal value of the fraction of epithelial cells, $\hat{x}$. Therefore, reaching the value of $\hat{x}$ is impossible due to the stochastic time evolution of the fraction of epithelial cells in the tumor. Instead, the optimization of phenotype switching rates provide the probability density function of the fraction of epithelial cells, such that the average fraction of epithelial cells is closest to the optimal value $\hat{x}$.

---


[1] Werner Horsthemke and René Lefever. *Noise-induced transitions: theory and applications in physics, chemistry, and biology.* Springer, 1984.

[2] Abhishek Dhar, Anupam Kundu, Satya N Majumdar, Sanjib Sabhapandit, and Grégory Schehr. Run-and-tumble particle in one-dimensional confining potentials: Steady-state, relaxation, and first-passage properties. *Physical Review E*, 99(3):032132, 2019.

[3] José M Sancho. External dichotomous noise: The problem of the mean-first-passage time. *Physical Review A*, 31(5):3523, 1985.

[4] Mathis Guéneau and Léo Touzo. Relating absorbing and hard wall boundary conditions for a one-dimensional run-and-tumble particle. *Journal of Physics A: Mathematical and Theoretical*, 57(22): 225005, 2024.

[5] Peter Häunggi and Peter Jung. Colored noise in dynamical systems. *Advances in chemical physics*,

89:239–326, 1994.

[6] Mathis Guéneau, Satya N Majumdar, and Grégory Schehr. Optimal mean first-passage time of a run-and-tumble particle in a class of one-dimensional confining potentials. *Europhysics Letters*, 145 (6):61002, 2024.
